# Patterns of sexual variation in hominoid mandibular morphology: a framework for interpreting the hominin fossil record

**DOI:** 10.1101/2022.06.15.496279

**Authors:** Lucía Nadal, Marta Mirazón Lahr

**Affiliations:** Leverhulme Centre for Human Evolutionary Studies, Department of Archaeology, University of Cambridge, Cambridge, UK; Turkana Basin Institute, Nairobi, Kenya

**Keywords:** Hominoids, mandible, sexual dimorphism, morphological variation, hominin diversity

## Abstract

For many species, sexual dimorphism is one of the major sources of intraspecific variation. This is the case in some extant great apes, such as gorillas and orangutans, and to a lesser degree in humans, chimpanzees and bonobos. This variation has been well documented in various aspects of these species skeletal anatomy, including differences in the size and shape of the body, cranium, canines, and cresting of males and females, but less is known about sexually dimorphic variation of great ape mandibles. This is particularly important for building robust analog models to interpreting variation in the early hominin fossil record which preserves a large proportion of isolated mandibles and partial mandibles. Here we describe the phenotypical expression of sexual dimorphism in the mandible of six extant hominoid species, including humans, using geometric morphometrics. Our analyses show that the extent of sexual dimorphism in mandibular size and shape amongst the species studied is not the same, as well as the presence of significant differences in the degree of sexual dimorphism being expressed at different sections of the mandible. Furthermore, we find significant differences in how sexual dimorphism is expressed phenotypically even amongst closely related species with small divergence times. We discuss the potential pathways leading to such variation and the implications for extinct hominin variability.

## 1. Introduction

Sexual dimorphism is an important source of intraspecific morphological variability among many primate species, including all extant great apes. The degree and particular expression of sexual dimorphism in living great apes has been the focus of multiple studies (e.g., Ashton, 1957; Schultz, 1962; Wood, 1975; Shea, 1983; Oxnard, 1987), particularly in relation to the relative magnitude and nature of intra-and inter-specific variation in cranial sexual dimorphism (e.g. O’Higgins et al., 1990; Berge and Penin, 2004; Schaefer et al., 2004; Cobb and O’Higgins, 2007; Balolia et al., 2013). These studies clearly show that the magnitude of sexual dimorphism amongst extant great apes differs, as does its anatomical expression. How these patterns of variation are expressed in other skeletal elements, including the mandible, has been less explored.

Mandibles and mandibular fragments account for a large percentage of the known hominin fossil record and are therefore a critical skeletal element to assess functional and taxonomic hominin variability. Extant hominoid primates, including humans, are commonly used as analogs to delimit this extinct variability (e.g. Quinney and Collard, 1995; Richmond and Jungers, 1995; Lockwood et al., 1996; Rosas et al., 2002; Ritzman et al., 2016). Yet, since there are differences in the extent and form of sexual dimorphism in living great apes, identifying how these differences are expressed in the mandible is critical for establishing analog boundaries of variation to inform palaeontological interpretations. Furthermore, the fragmentary nature of this record requires understanding the informative power that different aspects of the mandible might provide.

As expected, the shape of the mandible of different hominoids has been found to reflect functional (diet) and developmental (patterns of early growth) differences (Taylor, 2002, 2003, 2004, 2005), with sexual dimorphism the third significant source of variation. This is observed in most primates, among whom inter-specific differences in the expression of sexual dimorphism also show phylogenetic constraints, whereby closely related species express sexually dimorphic variation in similar ways (Plavcan, 2002). Yet, the few studies that focus on hominoid mandibles suggest that this is not the case in this group. Based on linear measurements, Wood et al. (1991) found that, in terms of magnitude of sexual dimorphism, *Gorilla* mandibles are closer to those of *Pongo* than to the less dimorphic, but more closely related, *Homo* and *Pan*. These results are consistent with those of Taylor (2006), who using ratios computed from linear measurements, found that species-specific patterns of sexual dimorphism in the hominoid mandible are mainly expressed in size, rather than shape differences. However, whether the differences observed in the mandible of males and females of each extant hominoid species and sub-species also reflect shape differences remains debated (Schmittbuhl et al., 2007). This raises two main issues – first, the roles of size and shape in defining the differences between male and female ape mandibles, and second, the taxonomic constraints on the expression of such differences.

The present study addresses these. We use geometric morphometric methods to explore interspecific patterns of size and shape sexual dimorphism in a sample of extant non-human great ape mandibles of known sex and in a sample of osteologically sexed modern human mandibles. We quantify and visualize shape differences in order to describe the observed phenotypic patterns and measure morphological disparities. By exploring the magnitude and particular expression of sexual dimorphism in the mandible of extant great apes, we aim at developing more robust comparative parameters for interpreting the morphological variability encountered in the hominin fossil record.

## 2. Methods

### 2.1 Samples

Non-human great ape mandibles included in this study are part of the Smithsonian Institution National Museum of Natural History Collection, and 3D models were accessed on the MorphoSource.org virtual repository (Project: Smithsonian Open Access, ID: 00000C955). All specimens are of known-sex and wild-shot, belonging to five species of extant apes (*Gorilla beringei* (n=12), *Gorilla gorilla* (n=32), *Pan troglodytes* (n=17), *Pongo pygmaeus* (n=10) and *Pongo abelii* (n=12)). See Supplementary Online Material (SOM) Table S1 for media accession numbers.

The modern human sample used consists of 21 mandibles from recent populations curated at the Duckworth Laboratory, University of Cambridge. Modern human 3D models were generated from Computer Tomography (CT) scans obtained at Addenbrooke’s Hospital using the InVesalius software (Version 3.1, Centre for Information Technology Renato Archer). The sex of these individuals is not known and was estimated on the basis of standard osteometric criteria in both mandible and associated cranium (Buikstra and Ubelaker, 1994). See SOM Table S1 for information on the human sample used.

All 3D models were standardized with a mesh decimation of 100,000 target faces in Meshlab (Cigoni et al., 2008), and the right hemimandible was extracted by bisecting meshes at the symphysis in Blender 3D (Version 2.91, Blender Foundation). Specimens included in this study exhibit a fully erupted M_3_, and no individuals showing significant damage or pathological morphologies were included in the analysis. Table 1 shows the complete sampled taxa and the collections that curate them.

**Table 1.** Sampled taxa

| Species | Male | Female | Total | Collection |
| --- | --- | --- | --- | --- |
| *Homo sapiens* | 14 | 7 | 21 | 1 |
| *Gorilla gorilla* | 24 | 8 | 32 | 2 |
| *Gorilla beringei* | 8 | 4 | 12 | 2 |
| *Pan troglodytes* | 8 | 9 | 17 | 2 |
| *Pongo abelii* | 6 | 6 | 12 | 2 |
| *Pongo pygmaeus* | 3 | 7 | 10 | 2 |
| **Total** | 63 | 41 | 104 |  |

\*Collection codes: 1=Duckworth Laboratory, University of Cambridge, 2= Smithsonian Institution National Museum of Natural History Collection.

### 2.2 Geometric Morphometrics

Landmark atlas design consisted of 20 fixed landmarks, 50 sliding semi-landmarks on equidistant curves and 331 surface semi-landmarks, placed on the right hemimandible (Figure 1). Detailed fixed and curved landmark definitions are shown in SOM Table S2.

**Figure 1.**
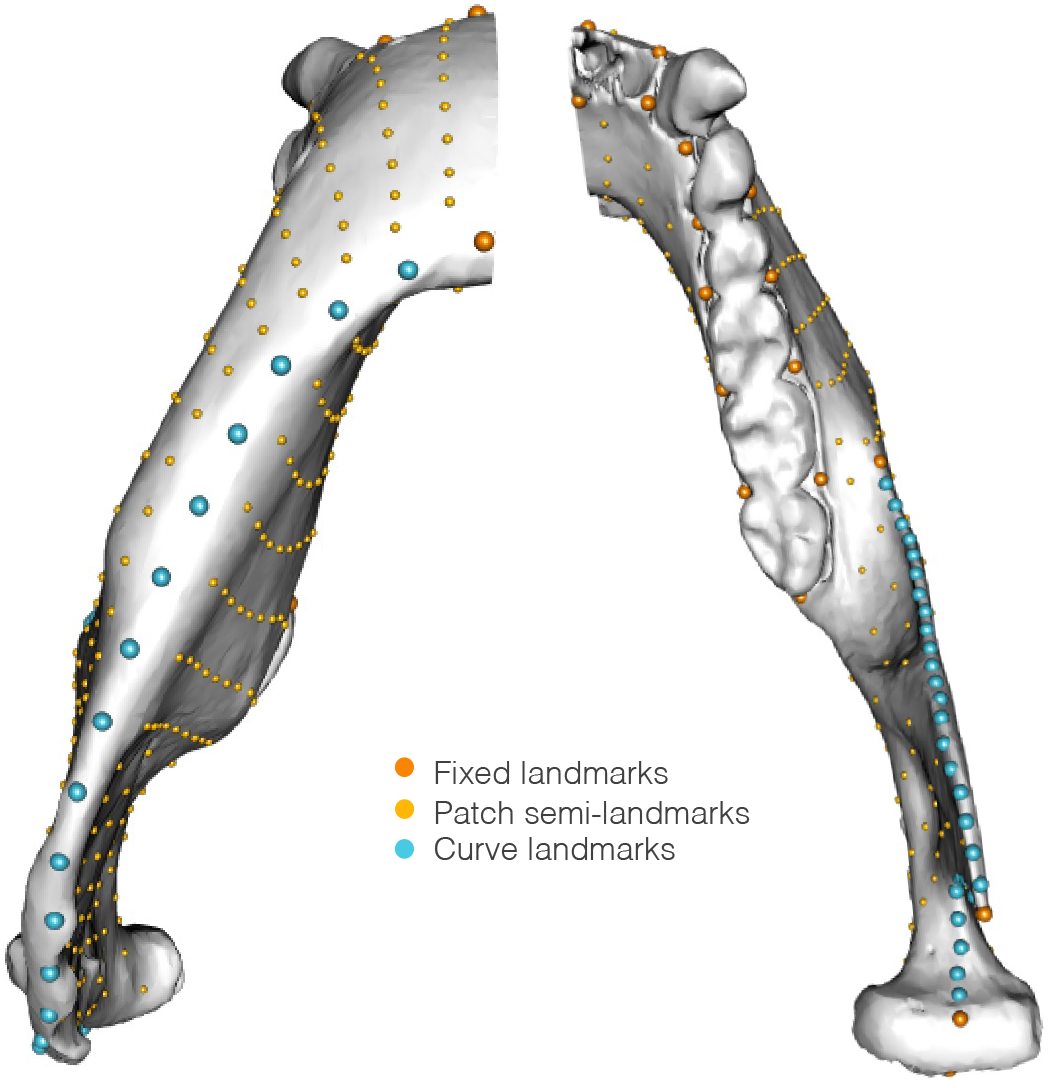
Landmark design showing fixed landmarks and sliding semi-landmarks on equidistant curves and surface semi-landmarks.

The Stratovan Checkpoint software (Version 2018.08.07, Stratovan Corporation) was used to obtain 3D landmark coordinate data. Further landmark patching, sliding by minimizing bending energy, and superimposition by Generalized Procrustes Analysis was done using the Morpho package in R (Schlager, 2017). This last step was carried out using the procSym() function, which also extracted the centroid size of each specimen. Centroid size was used in further analyses as a general measure of size.

### 2.3 Analysis

We calculated two measures of morphological disparity between males and females of each species. First, an overall measure of sexual shape dimorphism was calculated as the Procrustes distance between the mean female and mean male shapes for each species; second, a sexual size dimorphism index was calculated for each species as mean male centroid size divided by mean female centroid size, following Gibbons and Lovich (1990). Differences between species for both of these measurements were evaluated with a pairwise comparison. Additionally, overall intraspecific morphological disparity, measured as the Procrustes variance, and a corresponding post-hoc pairwise comparison was calculated for each species using the ‘morphol.disparity’ function from the package geomorph in R (Adams et al., 2021; Baken et al., 2021).

In order to further describe the nature of mandibular shape sexual dimorphism, a phenotypic trajectory analysis (PTA; Adams and Collyer 2009; Collyer and Adams 2013) was used to compare the magnitude and direction of shape change associated to sexual dimorphism between species, focusing on the inner and outer faces of both the mandibular ramus and corpus. These analyses were based on multifactorial linear models to quantify the variation in shape (i.e. aligned landmark coordinates) attributable to species and sex, with subsequent principal component analyses being conducted on the resulting fitted values, and least square (LS) means being used as trajectory endpoints. This method allows for the dimorphic shape change vectors to be projected onto the multidimensional morphospace, using male and female mean points for each species. A post-hoc pairwise comparison of the trajectories’ geometric attributes (path length and direction) was done using the ‘trajectory.analysis’ function in the RRPP package in R (Collyer and Adams, 2019).

The relationship between size and shape, and its sexually dimorphic expression, was investigated by calculating common allometric components using the CAC() function in the Morpho package in R (Schlager, 2017). Static allometric trajectories of males and females were obtained for each species by regressing the common allometric component to the logarithmic centroid size. Homogeneity of slopes was tested using Procrustes ANOVA in order to describe parallel or non-parallel static allometric trajectories between males and females of each species.

In order to visually maximize between-sex differences and highlight shape changes associated with this dimorphism, visualizations were generated by extracting the maximum male and female configurations emanating from a canonical variate analysis (CVA) carried out for each species and deforming the specimen closest to the species sample mean onto the coordinate configurations for each sex. Visualizations using mean male and female shapes for each species were also produced in order to corroborate the results obtained from maximum configurations.

Finally, in order to provide phenotypic dimensionality to the differences observed between males and females of each species, pairs of landmarks were chosen, and linear Euclidean distances were obtained to reflect linear measurements of symphyseal height, corpus width at M_2_-M_3_, symphysis base to coronoid process, symphysis base to condyle, and mandibular notch length (see SOM Table S3 and Figure S1 for the specific landmark boundaries and definitions of these measurements). Computation of these distances was done using the function ‘interlmkdist’ from the geomorph package in R (Adams et al., 2021; Baken et al., 2021). Assessment of the differences in these measurements between males and females was done using a student t-Test for each species.

## 3. Results

Figures 2A,B show the observed sexual dimorphism in mandibular shape (Procrustes distance between sexes) and size (sexual size dimorphism index) across the species sampled. *P. pygmaeus* displays the greatest differences in both the shape and size of male and female mandibles across all extant great apes studied, and the only statistically significant differences found with other extant hominoids (with *H. sapiens* in the case of shape: Z = 1.665, *p-value* = 0.044; and with *P. troglodytes* in the case of size: Z= 1.508, *p-value* = 0.047).

**Figure 2.**
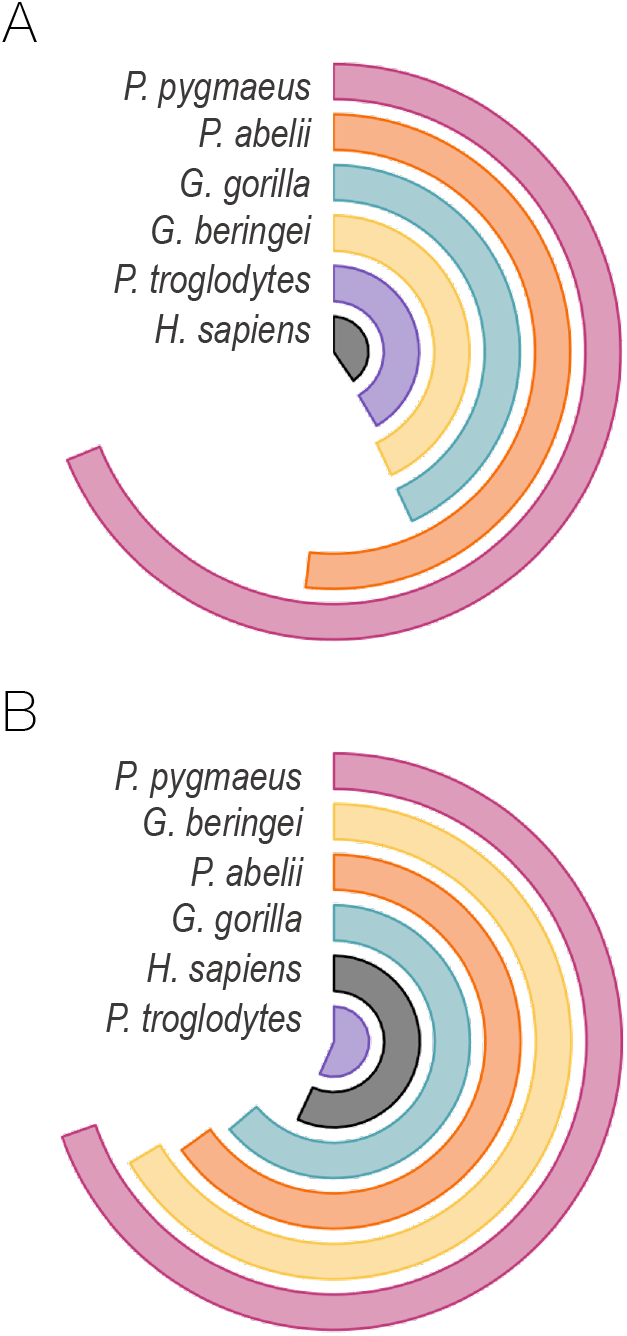
Radial bar charts of Sexual Shape Dimorphism measured as Procrustes distances (A), and Sexual Size Dimorphism Index (B), ranked from larger to smaller.

Among the other species studied, the relative extent of sexual dimorphism in the shape and size of the mandible differs. *P. abelii*, for example, shows higher values of shape dimorphism than African apes (both species of *Pongo* show substantially higher values than the rest of the species used in the study), but smaller size dimorphism than *G. beringei*. Similarly, while *H. sapiens* shows the smallest shape sexual dimorphism, it is in *P. troglodytes* mandibles that we observe the smallest differences in the size of male and female mandibles. Table 2 shows the exact species-specific distance values of size and shape differences between males and females ranked from smallest to largest.

**Table 2.** Sexual dimorphism in mandibular shape and size

| Species | Shape Dimorphism |  | Size Dimorphism Index |
| --- | --- | --- | --- |
| *Homo sapiens* | 0.02414522 | *Pan troglodytes* | 1.017416 |
| *Pan troglodytes* | 0.02492518 | *Homo sapiens* | 1.024081 |
| *Gorilla beringei* | 0.02576355 | *Gorilla gorilla* | 1.149577 |
| *Gorilla gorilla* | 0.02583594 | *Pongo abelii* | 1.172872 |
| *Pongo abelii* | 0.03110402 | *Gorilla beringei* | 1.196461 |
| *Pongo pygmaeus* | 0.04128420 | *Pongo pygmaeus* | 1.253491 |

\*Shape dimorphism given by Procrustes distance between sexes. Size Dimorphism Index obtained using Centroid size and following Gibbons & Lovich (1990).

Principal component and phenotypic trajectory analyses were conducted for the inner and outer faces of both corpus and ramus, with morphological disparities calculated for each species. Figure 3 shows the first and second principal components, which capture the main trends in morphological variation, and the projected trajectories of sexual dimorphism for each species, showing mean female and mean male points. The first principal component of the four mandibular sections analyzed captures the distinctiveness of *H. sapiens* in relation to all other non-human ape samples, of which the outer corpus reflects the largest differentiation. Procrustes ANOVA confirms the presence of significant variation in shape between species for all four mandibular sections, and post-hoc pairwise comparisons show *H. sapiens* to differ significantly from all other species in the inner ramus and both sections of the corpus (see SOM Table 5 for complete Procrustes ANOVA and pairwise results).

**Figure 3.**
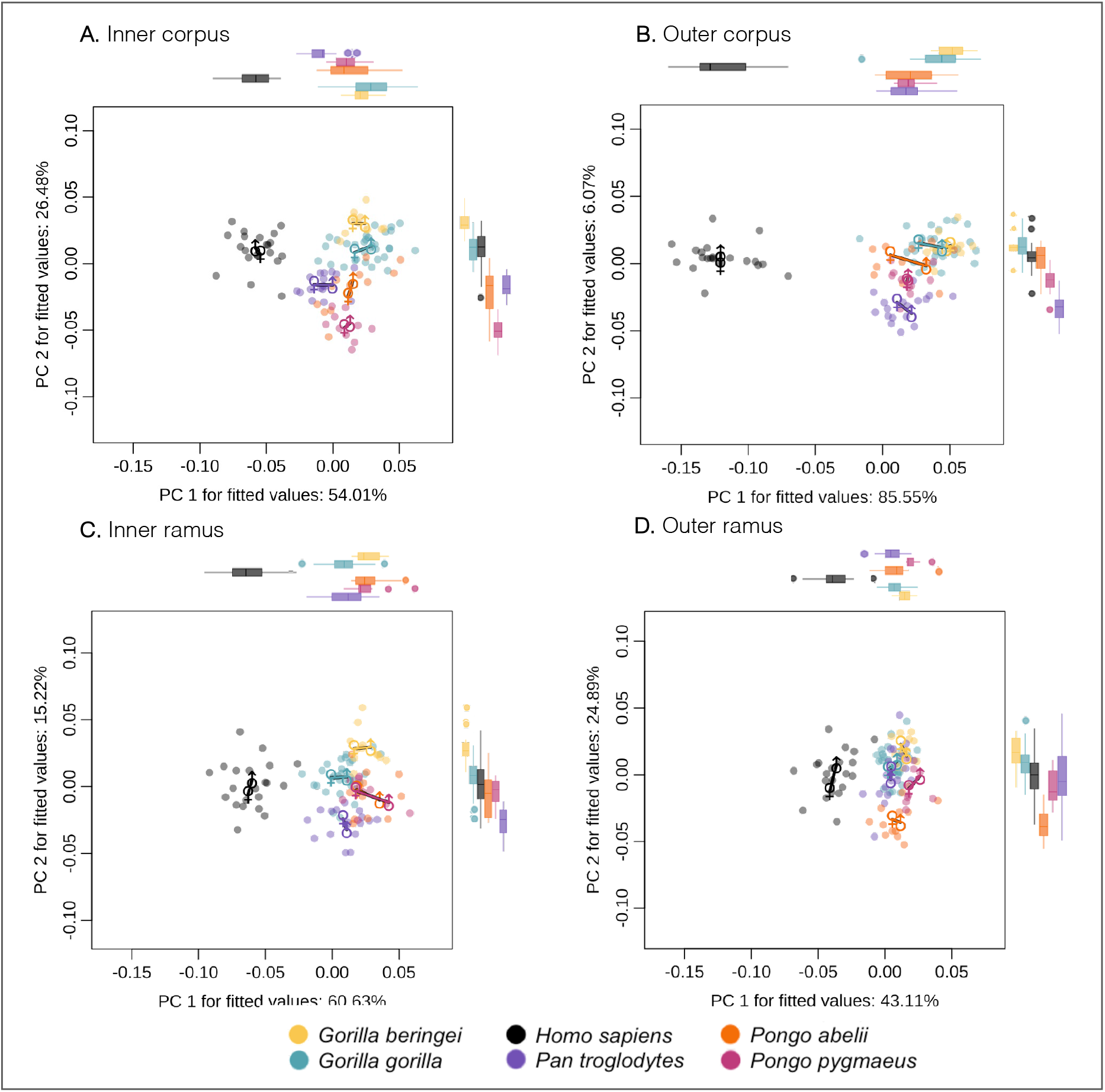
Shape dimorphism trajectories for each species projected onto the first two principal components showing male to female paths for inner corpus (A); inner ramus (B); outer corpus (C); and outer ramus (D) for *G. gorilla, G. beringei, H. sapiens, P. troglodytes, P. abelii* and *P. pygmaeus*. Marginal boxplots show distributions for each species at individual principal components.

Furthermore, the variability of *H. sapiens* on this first principal component tends to be relatively large when compared to all other species. This is further observed in the Procrustes variances obtained for *H. sapiens* which show the highest values obtained for all four mandibular sections. Notably, a corresponding post-hoc pairwise comparison shows these Procrustes variances to be significantly higher in *H. sapiens* when compared to all other species for the outer corpus and the inner ramus (See SOM Table S6 for complete Procrustes variances and corresponding post-hoc analyses).

In comparison, the second principal component captures some of the interspecific differences amongst the non-human great ape species studied. Of these, the inner corpus and the outer ramus sections of the mandible show the largest distances between *Gorilla* and *Pongo* samples, while the outer corpus and the inner ramus exhibit considerable differences between *Gorilla* and *Pan*. A post-hoc pairwise comparison reveals *G. beringei* and *P. pygmaeus* to be significantly different from every other species in both corpus sections, while *G. gorilla* differs significantly from all other species in both ramus sections. *P. troglodytes* is only significantly different from all other species in the outer ramus, and finally, *P. abelii* presents the lowest number of interspecific significant differences (see SOM Table 5 for complete Procrustes ANOVA and pairwise results).

Sexual dimorphism trajectories were projected onto the morphospace as paths from mean female and male points (Figure 3). A statistical evaluation of the differences in the magnitude (path length) and direction (path angle) of sexual dimorphism between species was done using pairwise comparisons on the dimorphism trajectories. In the case of magnitude, significant differences were only found in the inner and outer ramus. Of these, we observe significant differences in the magnitude of sexual dimorphism of the inner ramus between *P. pygmaeus* compared to *P. troglodytes* and both species of *Gorilla*, and of the outer between modern humans compared to *P. troglodytes* and *G. gorilla*. In contrast, significant differences in the direction of the dimorphism trajectories were found in all four sections of the mandible studied, all of which involve comparisons between *H. sapiens* and other species. The most extensive amongst these differences are observed between *H. sapiens* and *P. troglodytes* in all sections of the mandible. Further significant variation in the direction of dimorphism was found between modern humans and both species of *Gorilla* in the inner and outer corpus, both species of *Pongo* in the inner ramus, and *G. beringei* and *P. abelii*, respectively, in the outer ramus (see SOM Tables 7 and 8 for complete pairwise results).

Common allometric component scores (Mitteroecker et al. 2004) were calculated, and static allometric trajectories were obtained by regressing scores of males and females of each species to logarithmic centroid size; the homogeneity of male and female slopes was tested using Procrustes ANOVA. While no statistically significant differences were found between male and female static allometric trajectories for any of the species considered, it is interesting to note that the lowest associated p-values were found in *G. gorilla* (p-value= 0.075) and *P. pygmaeus* (p-value= 0.055). Of these two species, *G. gorilla* females exhibit steeper slopes in comparison to males, denoting higher allometric shape scores in females than those predicted for their size. In comparison, *P. pygmaeus* exhibits the opposite pattern, with males showing steeper slopes and higher allometric shape scores than those predicted for their size. Contrastingly, higher associated p-values are found for *P. abelii, G. beringei, P. troglodytes* and *H. sapiens* (Figure 4), depicting a more evident pattern of isometric dimorphism.

**Figure 4.**
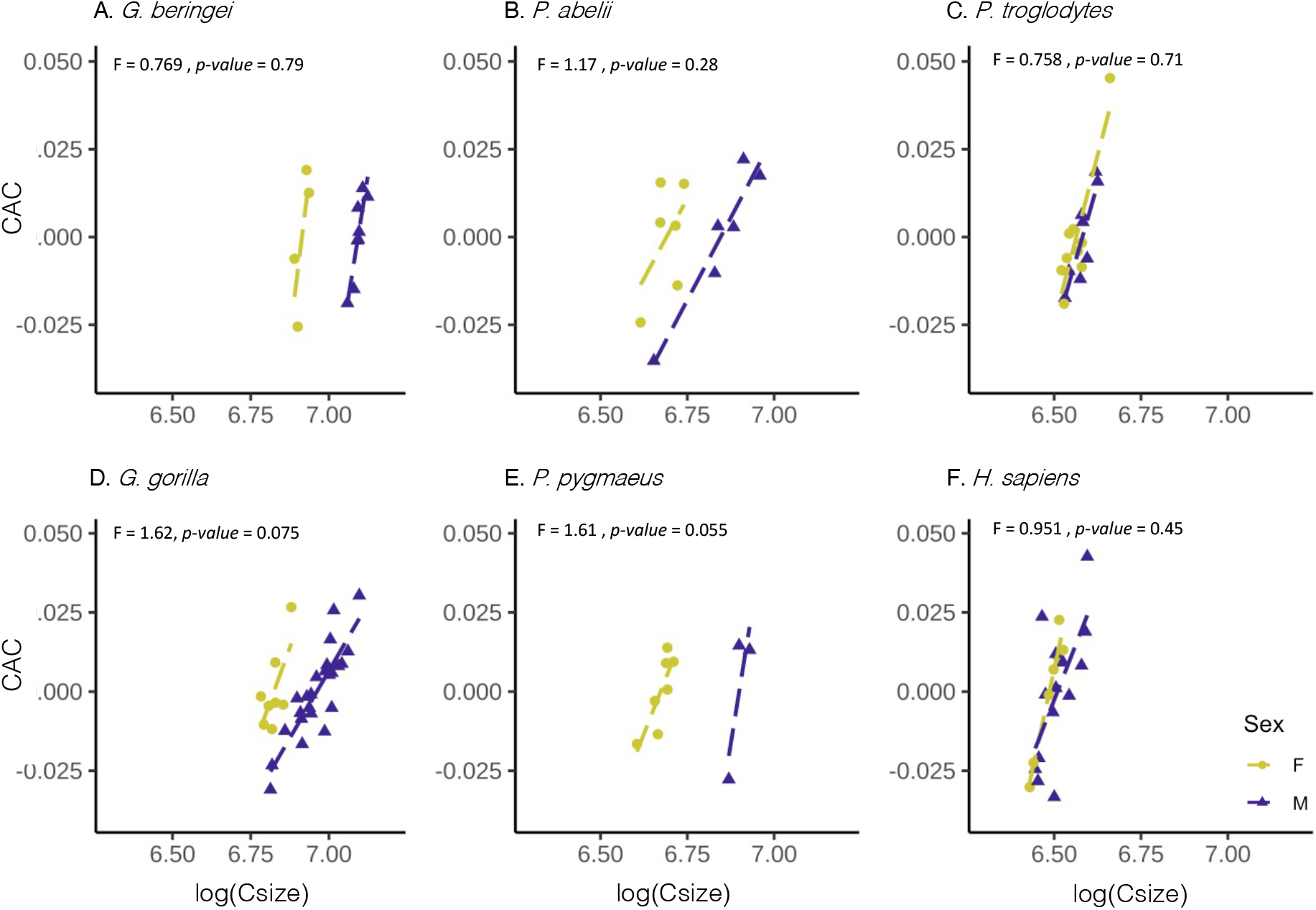
Static allometric trajectories showing regressions between logarithmic centroid size and the common allometric component (CAC) for males and females of *G. beringei* (A), *P. abelii* (B), *P. troglodytes* (C), *G. gorilla* (D), *P. pygmaeus* (E), and *H. sapiens* (F). Procrustes ANOVA results for individual species are shown

Canonical variate analyses and thin plate spline deformations were used in order to identify those aspects of morphology that best characterize the male and female mandibular shape in each of the six hominoid species being studied, as well as to graphically visualize these dimorphic differences by anchoring the produced male and female shapes at the symphysis (Figure 5). Among the differences observed, particularly notable is the positioning of the ramus on the coronal and sagittal planes. In the case of the coronal plane, females exhibit a posterior extension of the ramus with respect to males in *G. beringei, G. gorilla* and *P. pygmaeus*. Contrastingly, males exhibit a posterior extension of the ramus with respect to females in *P. abelii, P. troglodytes* and *H. sapiens*. In the sagittal plane, both species of *Pongo* and *Gorilla* share a dimorphic pattern in which females exhibit a more medial positioning of the ramus with respect to males. On the contrary, female rami are positioned laterally with respect to males only in *P. troglodytes* and, to a lesser degree, in *H. sapiens*.

**Figure 5.**
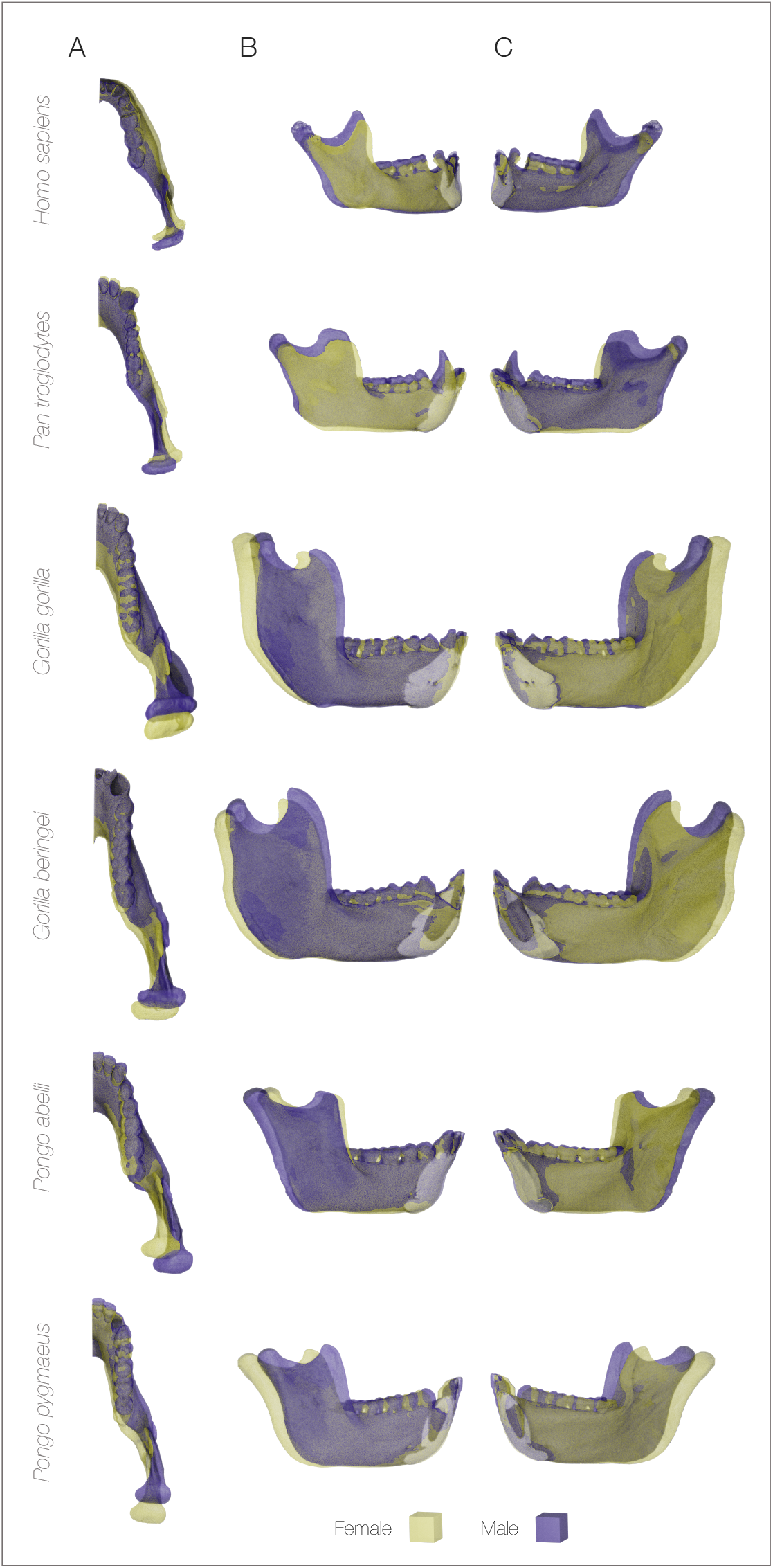
Overlapped visualizations obtained by canonical variate analysis, for maximum male and female shapes of each species. Shapes are aligned and anchored at the symphysis.

Finally, in order to add dimensionality to the male and female shape phenotypes, we computed linear distances between landmarks that reflect measurements of symphyseal height, corpus width at M_2_-M_3_, symphysis base to coronoid process length, symphysis base to condyle length, and mandibular notch length. A Student’s t-test (Table 3) revealed no significant differences for any of these measurements between males and females of either *H. sapiens* or *P. troglodytes*. Contrastingly, significant differences exist in all linear measurements computed, except corpus width, between males and females of both *Gorilla* species. In the case of *P. pygmaeus*, significant differences are found in all measurements except for mandibular notch, while *P. abelii* shows significant differences in all measurements except symphysis height. These results further confirm that chimpanzee and human male and female mandibles do not differ in shape, while the pronounced differences in the size of male and female mandibles of gorillas and orangutans are also expressed in particular shape differences.

**Table 3.** Student’s t-Test results from linear measurements of males and females of each species

| *G. beringei* | t | df | p-value |
| --- | --- | --- | --- |
| **Symphysis height** | **7.476** | **6.5236** | **0.000198** |
| Corpus width | 1.6587 | 5.5696 | 0.1521 |
| **Coronoid projection** | **9.8728** | **9.9408** | **1.87<sup>-6</sup>** |
| **Condyle projection** | **13.646** | **8.9211** | **2.79<sup>-7</sup>** |
| **Mandibular notch** | **2.5037** | **9.9649** | **3.13<sup>-2</sup>** |
| *G. gorilla* | t | df | p-value |
| **Symphysis height** | **3.8248** | **10.432** | **0.003101** |
| Corpus width | 1.9624 | 11.208 | 0.07502 |
| **Coronoid projection** | **6.5812** | **27.887** | **3.97<sup>-7</sup>** |
| **Condyle projection** | **6.1457** | **17.212** | **1.02<sup>-5</sup>** |
| **Mandibular notch** | **2.59** | **9.5** | **2.8<sup>-2</sup>** |
| *P. pygmaeus* | t | df | p-value |
| **Symphysis height** | **3.7235** | **2.4345** | **0.04775** |
| **Corpus width** | **4.4752** | **2.5413** | **0.0293** |
| **Coronoid projection** | **6.6143** | **3.0301** | **0.006822** |
| **Condyle projection** | **6.981** | **3.037** | **0.005796** |
| Mandibular notch | 1.9165 | 2.4023 | 0.174 |
| *P. abelii* | t | df | p-value |
| Symphysis height | 2.0823 | 5.7032 | 0.08489 |
| **Corpus width** | **4.837** | **9.9949** | **0.000686** |
| **Coronoid projection** | **3.7654** | **7.3652** | **0.006398** |
| **Condyle projection** | **3.2473** | **6.7175** | **0.01494** |
| **Mandibular notch** | **2.5456** | **7.4293** | **0.0365** |
| *H. sapiens* | t | df | p-value |
| Symphysis height | 0.39008 | 7.1298 | 0.7079 |
| Corpus width | 2.1932 | 10.083 | 0.05284 |
| Coronoid projection | 1.705 | 12.034 | 0.1138 |
| Condyle projection | 0.92569 | 17.197 | 0.3674 |
| Mandibular notch | -0.43946 | 15.612 | 0.666 |
| *P. troglodytes* | t | df | p-value |
| Symphysis height | 2.1403 | 11.427 | 0.05467 |
| Corpus width | -1.6294 | 13.309 | 0.1267 |
| Coronoid projection | -0.33183 | 12.578 | 0.7455 |
| Condyle projection | -0.6314 | 14.767 | 0.5374 |
| Mandibular notch | -0.82881 | 14.826 | 0.42 |

## 4 Discussion

This study set out to investigate patterns of sexual dimorphism in the mandible of extant great apes by asking two particular questions: 1) what is the magnitude of male-female differences in the size and shape of the mandible of different extant hominoid species? and 2) is mandibular sexual dimorphism expressed similarly in different extant hominoids? The results of the analyses above show a complex pattern of species-specific sexual dimorphism, which we discuss below.

### 4.1 Magnitude of sexual dimorphism in the size and shape of hominoid mandibles

Previous studies have highlighted the different features in which sexual dimorphism is expressed in ape species, particularly body size, canine size and cranial size/shape differences (e.g., Leutenegger and Cheverud, 1981; Kelley, 1995; Leigh, 1995a,b; Leigh and Shea, 1995; Schaefer, 2004; Cobb and O’Higgins, 2007), characterizing and quantifying the patterns that are clearly visible in wild and captive populations. From these observations, the species considered in the present study can be separated into two groups – the very sexually dimorphic gorillas and orangutans who are socially organized around intense single male dominance and competition, and the moderately dimorphic chimpanzees and humans who characteristically live in multi-male, multi-female societies (Plavcan, 2012). Ranking by the magnitude of size sexual dimorphism shows the extent to which different phenotypic traits are dimorphic. In terms of body size (measured as the male to female ratio in average body weight), *G. gorilla* are the most sexually dimorphic (2.37), followed by *P. pygmaeus* (2.23), *P. abelii* (2.23), *G. beringei graueri* (2.19*), *G. beringei beringei* (1.63*), *P. paniscus* (1.36), *P. troglodytes schweinfurthii* (1.29), and *H. sapiens* (1.15) (Leigh and Shea, 1995; *only one male data point for each subspecies of *G. beringei*). In terms of craniofacial size differences (measured as the difference between male and female centroid size), the ranking is similar, differing in the ordination of the less dimorphic group: most dimorphic crania are found in *G. gorilla*, followed by *P. pygmaeus, H. sapiens, P. troglodytes* and *P. paniscus* (Schaefer et al., 2004). As shown in Table 2, the pattern of size dimorphism in male and female ape mandibles (measured as mean male centroid size divided by mean female centroid size) that we obtain is different from both of these: the largest male-female differences in the mandible are observed in *P. pygmaeus* (1.253), followed by *G. beringei* (1.196), *P. abelii* (1.173), *G. gorilla* (1.149), *H. sapiens* (1.024) and *P. troglodytes* (1.017). This diversity in how sexual dimorphism is expressed morphologically in adults of different ape species is the outcome of private selective pressures on different phenotypes that, controlled by trade-offs between energetic ecology and maturation, shaped growth rates and duration of life-history phases (Leigh and Shea, 1996; Leigh, 2001; Pereira and Leigh, 2003; Dirks and Bowman, 2007). These private evolutionary histories resulted in differences in body, brain and dental development (Leigh, 1995 a,b; Kuzawa et al., 2014) that underscore each species’ pattern of male and female differences in size and shape of different anatomical parts.

We also analyzed the magnitude of shape dimorphism between male and female adult ape mandibles (measured as the Procrustes distance between the sexes). As is also observed in the craniofacial skeleton (Schaefer et al. 2004, Pitirri and Begun 2019), shape differences are more pronounced in orangutan mandibles [*P. pygmaeus* (0.041), *P. abelii* (0.031)] than in those of African apes [*G. gorilla* (0.026), *G. beringei* (0.026), *P. troglodytes* (0.025) and *H. sapiens* (0.024)]. This is consistent with studies that show that mandible shape has a strong phylogenetic signal (e.g. Schmittbuhl et al., 2007; Balolia et al., 2020), and highlights a strong integration between cranial and mandibular shape (Bastir and Rosas, 2005). Notably, all the ape species included in this study show relatively higher magnitudes of shape sexual dimorphism in the mandibular ramus than in the corpus. Musculo-skeletal development has been recognized as a significant contributor to mandibular sexual dimorphism in human (Rosas et al., 2002) and non-human primates (Plavcan, 2002), and to result in high levels of intra-specific variability (Wood and Lieberman, 2001). We attribute our finding of a shared hominoid pattern of greater shape dimorphism in the mandibular ramus than the corpus to differences in musculo-skeletal development between males and females, whereby, as anchor to key masticatory muscles, the mandibular ramus is integrated into broader cranio-mandibular anatomical complexes than the corpus. In this framework, the relatively greater shape sexual dimorphism of the ramus than the corpus would be associated to the development of a non-isometric sufficiently large surface area dedicated to muscle attachments in male apes, with the insertion for the masseter located on the outer face and the medial pterygoid insertion on the inner face. Both muscles are the main elevators of the mandible, suggesting that hominoids express a strong sexual dimorphism in bite force, a notion previously suggested by Demes and Creel (1988).

### 4.2 Species-specific patterns of shape differences in male and female ape mandibles

Our analyses reveal significant interspecific variability in the expression of mandibular sexual dimorphism, even amongst closely related ape species. This observation is consistent with those of previous studies, such as Schmittbuhl et al. (2007), who also describe species-specific degrees of sexual dimorphism in the overall mandibular outline of all extant hominoids. Our results allow us to further quantify and describe some of these ape species-specific dimorphism patterns.

This study included two species of orangutan, *P. pygmaeus* and *P. abelii*, known for their high levels of sexual dimorphism in size and in the outline shape of the mandible (Schmittbuhl et al., 2007). As expected, we also observe these differences, but extend them to the size-corrected shape of the male and female mandibles of these two species. Although we found no significant variation in the magnitude or the direction of the shape dimorphism, we observed notable differences in phenotypic expression and static allometric trajectories. The mandibular corpus of both *P. pygmaeus* and *P. abelii* is broader in males than females, but, once corrected for size differences, male *P. pygmaeus* mandibles are shorter than female ones, with a ramus that is more vertically oriented and more anteriorly positioned, with a relatively shorter posterior dentition. In contrast, in *P. abelii*, male and female mandibles have similar inferior corpus lengths, and it is the more posteriorly oriented rami in males that make their mandible relatively longer than female ones, with a slightly longer molar row. Schmittbuhl and colleagues propose both neutral and functional explanations for the mandibular polymorphism in size and outline shape they observe in *Pongo* (Schmittbuhl et al., 2007). They emphasize the role of drift following allopatric genetic isolation in shaping “distinct evolutionary trends” in *P. pygmaeus* and *P. abelii* (Schmittbuhl et al., 2007), sister species that split approximately 1 million years ago (Yousef et al. 2021). Yet, orangutans from Sumatra and Borneo show a number of adaptive differences (Nater et al. 2017, Mattle-Greminger et al. 2018), and we would argue that the very nature of phenotypic variability associated with sexual dimorphism entails a non-neutral evolutionary mode. Accordingly, Vogel and colleagues stress differences in dietary ecology between Sumatran and Borneo orangutans as the main factor explaining their different expression of sexual dimorphism (Vogel et al., 2014), while differences in the frequency and length of male/female associations, and accompanying levels of female stress as measured by fecal cortisol levels (Kunz et al., 2021), offer another socioecological factor shaping their different adaptive landscapes.

A second case for interspecific divergence of sexual dimorphism patterns in closely related species can be observed between eastern (*G. beringei*) and western gorillas (*G. gorilla*). A previous study found no differences in mandibular dimorphism between these species (Taylor, 2006), and indeed, male and female mandibles of both groups share a number of traits (broader male than female mandibles, similar male and female inferior mandibular length, more posteriorly placed rami in females resulting in overall longer female mandibles with slightly longer posterior dentition). However, our analyses also show that *G. gorilla* displays overall higher levels of shape mandibular dimorphism, as well as dimorphism in static allometric trajectories, than *G. beringei*, whereby smaller male and female mandibles are relatively more similar in shape to each other than larger ones. In contrast, *G. beringei* shows a pattern of isometric dimorphism, whereby differences between males and females are more strongly manifested in size than in shape (as can be seen by the isometrically larger male *G. beringei* rami in Figure 5), and male and female static allometric trajectories are parallel. These interspecific variations are consistent with previous analyses of body size dimorphism in these species which show disparities in sex-differentiated growth patterns between eastern and western gorillas, resulting in *G. beringei* expressing a higher degree of body size dimorphism than *G. gorilla* (Taylor, 1997). These distinct growth patterns are directly related to the differences in life-history traits that have been documented between both species, with western gorillas exhibiting relatively slower physical maturation patterns than eastern gorillas (Breuer et al., 2009; McFarlin et al., 2013; Galvany et al., 2017; Kralick et al., 2017). Dietary ecology has been proposed as the main explanation for these differences, where the seasonality constraints imposed by frugivory in western gorillas result in reduced energy allocated to physical maturation (Doran and McNeilage, 2001). Other ecological factors, such as foraging range, predation risk and degree of arboreality, have also been suggested to play a role in shaping the differences in life-history traits of gorillas (Breuer et al., 2009).

The lowest levels of sexual dimorphism in both shape and size were found in *H. sapiens* and *P. troglodytes*, in agreement with previous findings (Wood et al., 1991). Nonetheless, our results show a remarkable divergence in dimorphism patterns between *H. sapiens* and *P. troglodytes* in terms of significantly different expression of shape dimorphism of the inner and outer faces of both corpus and ramus. This discrepancy in sexual dimorphism patterns between humans and chimpanzees has been previously observed in linear measurements and explained as being the result of differences in life-history traits that affect processes of bone resorption and deposition in the mandible (Humphrey et al., 1999). Even though both of these species exhibit relatively small mandibles in comparison to other extant apes, functional differences associated to their distinct diets are clearly reflected anatomically in the relative size of the anterior dentition and the height of the ramus (Humphrey et al., 1999). While human mandibles show a relatively tall ramus and small anterior dentition, the opposite pattern is true in chimpanzee mandible who engage their mouth and dentition in feeding behaviors such as stripping and peeling. These distinct morphologies have also been noted to have significantly different growth rates, with human mandibles displaying a unique growth pattern in which the ascending ramus attains adult dimensions earlier than any other ape, including chimpanzees (Humphrey, 1999). In conjunction, morphological differences related to dietary ecology and distinct growth patterns related to life-history traits have resulted in similar levels but significantly different patterns of sexual dimorphism in the mandibles of humans and chimpanzees.

### 4.3 Implications for the interpretation of the hominin fossil record

Mandibles and mandibular fragments represent a substantial proportion of the hominin fossil record and consequently play a significant role in interpretations of hominin taxonomy and evolutionary trends. Our results have implications for such interpretations.

Firstly, the magnitude of sexual dimorphism is not equally expressed across different sections of the mandible. Of particular note is the fact that mandibular rami are significantly more sexually dimorphic than the mandibular corpus. This suggests that, in terms of sexual dimorphism, the informative value of the preserved aspects of mandibular morphology should be considered when analyzing fossil hominin samples. Secondly, the interspecific variability in patterns of expression of sexual dimorphism observed even amongst closely related species of extant hominoids suggests that careful consideration should be given to the choice of extant analogs to infer variation due to sexual dimorphism in this fossil record. An appropriate comparative model should include a taxonomically variable extant model sample that reflects different morphological configurations of how sexual dimorphism may be expressed in the size and shape of ape mandibles. Lastly, the developmental processes that underlie species-specific patterns of sexual dimorphism point to differences in the dietary ecology of closely related species as playing a significant role in modifying the expression of mandibular sexual dimorphism. In the last decade, our understanding of the dietary ecology of extinct hominins has greatly increased, and a growing body of evidence points towards a wide range of dietary strategies (e.g., Cerling et al., 2013; Lüdecke et al., 2018). The results of this study suggest that such variability in dietary ecologies would have favored substantial diversity in patterns of sexual dimorphism amongst hominins.

## 5. Conclusions

Differences in the phenotypic expression of mandibular sexual dimorphism do not appear to be phylogenetically constrained, and closely related species of extant hominoids can exhibit divergent patterns of expression. Moreover, single highly integrated anatomical units, such as the mandible, might show variable degrees of sexual dimorphism in different sections. Two of the findings of this study are of particular significance for the interpretation of the variability encountered in the mandibular hominin fossil record. Firstly, changes in the pattern of expression of sexual dimorphism are observed even between species with short divergence times, cautioning against homogenizing the pattern of mandibular sexual dimorphism in extinct hominin species. Secondly, modern humans and chimpanzees show remarkably distinct dimorphic variation in mandibular morphology, questioning the use of *Pan*, or any other single hominoid species, as an analog to frame the morphological variability of fossil hominins. The complex and particular expression of sexual dimorphism found amongst different ape mandibles implies that no one species is an adequate analog for extinct hominins, and more inclusive comparative models that capture their diversity should be used.

## Acknowledgments

This study was funded by a CONACYT-Cambridge Trust doctoral scholarship, and the European Research Council Advanced Awards In-Africa (ERC #295907) and Ngipalajem (ERC #101020478). The authors would like to thank the Duckworth Laboratory, University of Cambridge for access to the modern human samples, as well as the Smithsonian Institution National Museum of Natural History Collection for access to the non-human 3D models through the virtual repository MorphoSource (Project: Smithsonian Open Access, ID: 00000C955).

## References

Adams, D.C., Collyer, M.L., 2009. A general framework for the analysis of phenotypic trajectories in evolutionary studies. Evolution. 63, 1143–1154.

Adams, D., Collyer, M., Kaliontzopoulou, A., Baken, E. 2021. Geomorph: Software for geometric morphometric analyses. R package version 4.0.2.

Ashton, E.H., 1957. Age changes in dimensional differences between the skulls of male and female apes. Proceedings of the Zoological Society of London. 128, 259–266.

Baken, E.K., Collyer, M.L., Kaliontzopoulou, A., Adams, D.C., 2021. geomorph v4.0 and gmShiny: Enhanced analytics and a new graphical interface for a comprehensive morphometric experience. Methods in Ecology and Evolution. 12, 2355–2363.

Balolia, K.L., Soligo, C., Lockwood, C.A., 2013. Sexual Dimorphism and Facial Growth Beyond Dental Maturity in Great Apes and Gibbons. International Journal of Primatology. 34, 361–387.

Bastir, M., Rosas, A., 2005. Hierarchical nature of morphological integration and modularity in the human posterior face. American Journal of Physical Anthropology. 128, 26–34.

Berge, C., Penin, X., 2004. Ontogenetic allometry, heterochrony, and interspecific differences in the skull of African apes, using tridimensional Procrustes analysis. American Journal of Physical Anthropology. 124, 124–138.

Breuer, T., Breuer-Ndoundou Hockemba, M., Olejniczak, C., Parnell, R. J., Stokes, E. J., 2009. Physical maturation, life-history classes and age estimates of free-ranging western gorillas - Insights from Mbeli Bai, Republic of Congo. American Journal of Primatology. 71, 106–119.

Buikstra, J.E., Ubelaker, D.H., 1994. Standards for Data Collection from Human Skeletal Remains. Fayetteville: Arkansas Archeological Survey Research Series No. 44.

Cerling, T.E., Manthi, F.K., Mbua, E.N., Leakey, L.N., Leakey, M.G., Leakey, R.E., Brown, F.H., Grine, F.E., Hart, J.A., Kaleme, P., Roche, H., Uno, K.T., Wood, B.A., Klein, R.G., 2013. Stable isotope-based diet reconstructions of Turkana Basin hominins. Proceedings of the National Academy of Sciences. 110, 10501–10506.

Cignoni, P., Corsini, M., Ranzuglia, G., 2008. Meshlab: an open-source 3d mesh processing system. ERCIM News, Vol. 73, p. 47–8.

Cobb, S.N., O’Higgins, P., 2007. The ontogeny of sexual dimorphism in the facial skeleton of the African apes. Journal of Human Evolution. 53, 176–190.

Collyer, M.L., Adams, D.C., n.d. Phenotypic Trajectory Analysis: Comparison of Shape Change Patterns in Evolution and Ecology. Hystrix, the Italian Journal of Mammalogy. 24, 75–83.

Collyer, M.L., Adams, D.C., 2019. RRPP: linear model evaluation with randomized residuals in a permutation procedure. R package version 0.4. 0.

Demes, B., Creel, N., 1988. Bite force, diet, and cranial morphology of fossil hominids. Journal of Human Evolution. 17, 657–670.

Dirks, W., Bowman, J.E., 2007. Life history theory and dental development in four species of catarrhine primates. Journal of Human Evolution. 53, 309–320.

Doran, D.M, McNeilage A., 2001. Subspecific variation in gorilla behavior: the influence of ecological and social factors. In: Robbins, MM., Sicotte, P., Stewart, KJ. (Eds.), Mountain gorillas: three decades of research at Karisoke. Cambridge: Cambridge University Press. pp 123–149.

Galbany, J., Abavandimwe, D., Vakiener, M., Eckardt, W., Mudakikwa, A., Ndagijimana, F., Stoinski, T.S., McFarlin, S.C., 2017. Body growth and life history in wild mountain gorillas (Gorilla beringei beringei) from Volcanoes National Park, Rwanda. American Journal of Physical Anthropology. 163, 570–590.

Gibbons, J.W., Lovich, J.E., 1990. Sexual Dimorphism in Turtles with Emphasis on the Slider Turtle (Trachemys scripta). Herpetological Monographs. 4, 1.

Humphrey, L.T. 1999. Relative mandibular growth in humans, gorillas and chimpanzees. In: Hoppa, R., Fitzgerald C. (Eds.), Human Growth in the Past: Studies from Bones and Teeth. Cambridge: Cambridge University Press, pp. 65–87.

Humphrey, L.T., Dean, M.C., Stringer, C.B., 1999. Morphological variation in great ape and modern human mandibles. Journal of Anatomy. 195, 491–513.

Kelley, J., 1995. Sexual dimorphism in canine shape among extant great apes. American Journal of Physical Anthropology. 96, 365–389.

Kralick, A. E., Loring Burgess, M., Glowacka, H., Arbenz-Smith, K., McGrath, K., Ruff, C. B., Chan, K. C., Cranfield, M. R., Stoinski, T. S., Bromage, T. G., Mudakikwa, A., McFarlin, S. C., 2017. A radiographic study of permanent molar development in wild Virunga mountain gorillas of known chronological age from Rwanda. American Journal of Physical Anthropology. 163, 129–147.

Kunz, J.A., Duvot, G.J., Noordwijk, M.A. van, Willems, E.P., Townsend, M., Mardianah, N., Atmoko, S.S.U., Vogel, E.R., Nugraha, T.P., Heistermann, M., Agil, M., Weingrill, T., Schaik, C.P. van, 2021. The cost of associating with males for Bornean and Sumatran female orangutans: a hidden form of sexual conflict? Behavioral Ecology and Sociobiology. 75, 6.

Kuzawa, C.W., Chugani, H.T., Grossman, L.I., Lipovich, L., Muzik, O., Hof, P.R., Wildman, D.E., Sherwood, C.C., Leonard, W.R., Lange, N., 2014. Metabolic costs and evolutionary implications of human brain development. Proceedings of the National Academy of Sciences. 111, 13010–13015.

Leigh, S.R., 1995a. Ontogeny and the evolution of body size dimorphism in primates. Anthropologie. 33, 17–28.

Leigh, S.R., 1995b. Socioecology and the ontogeny of sexual size dimorphism in anthropoid primates. American Journal of Physical Anthropology. 97, 339–356.

Leigh, S.R., 2001. Evolution of human growth. Evolutionary Anthropology: Issues, News, and Reviews. 10, 223–236.

Leigh, S.R., Shea, B.T., 1995. Ontogeny and the evolution of adult body size dimorphism in apes. American Journal of Primatology. 36, 37–60.

Leigh, S.R., Shea, B.T., 1996. Ontogeny of body size variation in African apes. American Journal of Physical Anthropology. 99, 43–65.

Leutenegger, W., Cheverud, J., 1981. Correlates of sexual dimorphism in primates: Ecological and size variables. International Journal of Primatology. 3, 387–402.

Lockwood, C.A., Richmond, B.G., Jungers, W.L., Kimbel, W.H., 1996. Randomization procedures and sexual dimorphism in Australopithecus afarensis. Journal of Human Evolution. 31, 537–548.

Lüdecke, T., Kullmer, O., Wacker, U., Sandrock, O., Fiebig, J., Schrenk, F., Mulch, A., 2018. Dietary versatility of early Pleistocene hominins. Proceedings of the National Academy of Sciences.115 (52), 13330–13335.

Mattle-Greminger, M.P., Sonay, T.B., Nater, A., Pybus, M., Desai, T., Valles, G. de, Casals, F., Scally, A., Bertranpetit, J., Marques-Bonet, T., Schaik, C.P. van, Anisimova, M., Krützen, M., 2018. Genomes reveal marked differences in the adaptive evolution between orangutan species. Genome Biology. 19, 193.

McFarlin, S.C., Barks, S.K., Tocheri, M.W., Massey, J.S., Eriksen, A.B., Fawsett, K.A., Stoinski, T.S., Hof, P.R., Bromage, T.G., Mudadidwa, A., Cranfield, M.R., Sherwood, C.C., 2013. Early brain growth cessation in wild Virunga mountain gorillas (Gorilla beringei beringei). American Journal of Primatology. 75, 450–463.

Mitteroecker, P., Gunz, P., Bernhard, M., Schaefer, K., Bookstein, F.L., 2004. Comparison of cranial ontogenetic trajectories among great apes and humans. Journal of Human Evolution. 46, 679–698.

Nater, A., Mattle-Greminger, M.P., Nurcahyo, A., Nowak, M.G., Manuel, M. de, Desai, T., Groves, C., Pybus, M., Sonay, T.B., Roos, C., Lameira, A.R., Wich, S.A., Askew, J., Davila-Ross, M., Fredriksson, G., Valles, G. de, Casals, F., Prado-Martinez, J., Goossens, B., Verschoor, E.J., Warren, K.S., Singleton, I., Marques, D.A., Pamungkas, J., Perwitasari-Farajallah, D., Rianti, P., Tuuga, A., Gut, I.G., Gut, M., Orozco-terWengel, P., Schaik, C.P. van, Bertranpetit, J., Anisimova, M., Scally, A., Marques-Bonet, T., Meijaard, E., Krützen, M., 2017. Morphometric, Behavioral, and Genomic Evidence for a New Orangutan Species. Current Biology. 27, 3487–3498.

O’Higgins, P., Moore, W.J., Johnson, D.R., McAndrew, T.J., Flinn, R.M., 1990. Patterns of cranial sexual dimorphism in certain groups of extant hominoids. Journal of Zoology. 222, 399–420.

Oxnard, C. E., 1987. Fossils. teeth and sex: New perspectives on human evolution. Seattle: University of Washington Press.

Pereira, M.E., Leigh, S.R., 2003. Modes of primate development. In: Kappeler, P.M., Pereira, M.E. (Eds.), Primate Life Histories and Socioecology. University of Chicago Press, Chicago, pp. 149–176.

Pitirri, M.K., Begun, D., 2019. A new method to quantify mandibular corpus shape in extant great apes and its potential application to the hominoid fossil record. American Journal of Physical Anthropology. 168, 318–328.

Plavcan, J.M., 2002. Taxonomic variation in the patterns of craniofacial dimorphism in primates. Journal of Human Evolution. 42, 579–608.

Plavcan, J.M., 2012. Sexual Size Dimorphism, Canine Dimorphism, and Male-Male Competition in Primates. Human Nature. 23, 45–67.

Quinney, P.S., Collard, M., 1995. Sexual Dimorphism in the Mandible of Homo neanderthalensis and Homo sapiens: Morphological Patterns and Behavioural Implications. In: Sinclair, A.G.M., Slater, E.A., Gowlett, J.A.J. (Eds.), Archaeological Sciences. Oxbow Books, pp. 420–425.

Richmond, B.G., Jungers, W.L., 1995. Size variation and sexual dimorphism in Australopithecus afarensis and living hominoids. Journal of Human Evolution. 29, 229–245.

Ritzman, T.B., Terhune, C.E., Gunz, P., Robinson, C.A., 2016. Mandibular ramus shape of Australopithecus sediba suggests a single variable species. Journal of Human Evolution. 100, 54–64.

Rosas, A., Bastir, M., Martínez-Maza, C., Castro, J.M.B. de, 2002. Sexual dimorphism in the Atapuerca-SH hominids: the evidence from the mandibles. Journal of Human Evolution. 42, 451–474.

Schaefer, K., Mitteroecker, P., Gunz, P., Bernhard, M., Bookstein, F.L., 2004. Craniofacial sexual dimorphism patterns and allometry among extant hominids. Annals of Anatomy - Anatomischer Anzeiger. 186, 471–478.

Schlager, S., 2017. Morpho and Rvcg – Shape Analysis in R. In: Zheng, G., Li, S., Szekely, G. (Eds.), Statistical Shape and Deformation Analysis. New York: Academic Press, pp. 217–256.

Schmittbuhl, M., Rieger, J., Minor, J.L., Schaaf, A., Guy, F., 2007. Variations of the mandibular shape in extant hominoids: Generic, specific, and subspecific quantification using elliptical fourier analysis in lateral view. American Journal of Physical Anthropology. 132, 119–131.

Schultz, A.H., 1962. Metric age changes and sex differences in primate skull. Zeitschrift für Morphologie und Anthropologie. 3, 239–255.

Shea, B.T., 1983. Allometry and heterochrony in the African apes. American Journal of Physical Anthropology. 62, 275–289.

Taylor, A.B., 1997. Relative growth, ontogeny, and sexual dimorphism in gorilla (Gorilla gorilla gorilla and G. g. beringei): Evolutionary and ecological considerations. American Journal of Primatology. 43, 1–31.

Taylor, A.B., 2006. Feeding behaviour, diet, and the functional consequences of jaw form in orangutans, with implications for the evolution of Pongo. Journal of Human Evolution. 50, 377–393.

Vogel, E.R., Zulfa, A., Hardus, M., Wich, S.A., Dominy, N.J., Taylor, A.B., 2014. Food mechanical properties, feeding ecology, and the mandibular morphology of wild orangutans. Journal of Human Evolution. 75, 110–124.

von Cramon-Taubadel, N., 2011. Global human mandibular variation reflects differences in agricultural and hunter-gatherer subsistence strategies. Proceedings of the National Academy of Sciences, 108, 19546–19551.

Yousaf, A., Liu, J., Ye, S., Chen, H., 2021. Current Progress in Evolutionary Comparative Genomics of Great Apes. Frontiers in Genetics. 12, 657468.

Wood, B. A., 1975. An analysis of sexual dimorphism in primates. PhD thesis, University of London.

Wood, B.A., Li, Y., Willoughby, C., 1991. Intraspecific variation and sexual dimorphism in cranial and dental variables among higher primates and their bearing on the hominid fossil record. Journal of Anatomy. 174, 185–205.

Wood, B., Lieberman, D.E., 2001. Craniodental variation in Paranthropus boisei: A developmental and functional perspective. American Journal of Physical Anthropology. 116, 13–25.

