## Supplementary Material for "Patterns of sexual variation in hominoid mandibular morphology: a framework for interpreting the hominin fossil record"

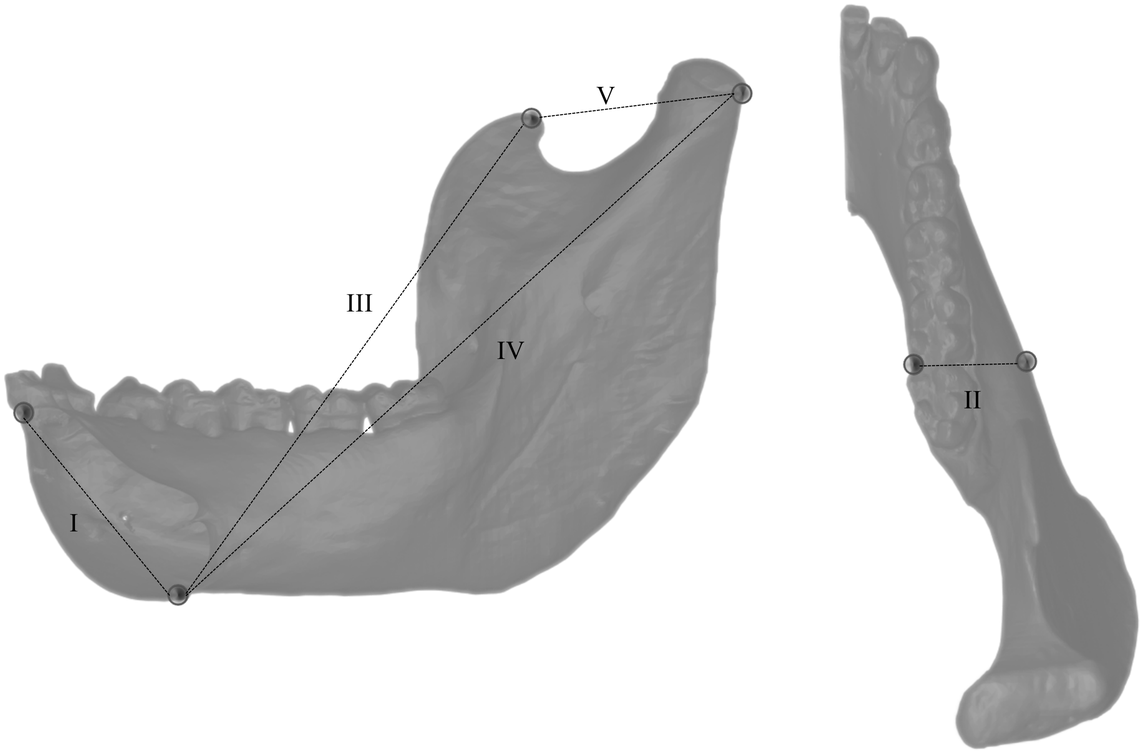


**SOM Figure S1.** Linear measurements extracted from landmarks. See Table S3 for definitions.

### SOM Table S1

Details of specimens included in the study

| # | Media accession number | Digital Repository | ID in Collection | Collection | Species | Sex |
|---|---|---|---|---|---|---|
| 1 | - | - | AF 115124 | Duckworth Laboratory, University of Cambridge | *Homo sapiens* | Male |
| 2 | - | - | DC 1735 | Duckworth Laboratory, University of Cambridge | *Homo sapiens* | Female |
| 3 | - | - | DC 1737 | Duckworth Laboratory, University of Cambridge | *Homo sapiens* | Male |
| 4 | - | - | DC 1738 | Duckworth Laboratory, University of Cambridge | *Homo sapiens* | Female |
| 5 | - | - | DC 1447 | Duckworth Laboratory, University of Cambridge | *Homo sapiens* | Male |
| 6 | - | - | AUS 027 | Duckworth Laboratory, University of Cambridge | *Homo sapiens* | Male |
| 7 | - | - | AUS 046 | Duckworth Laboratory, University of Cambridge | *Homo sapiens* | Female |
| 8 | - | - | AUS 047 | Duckworth Laboratory, University of Cambridge | *Homo sapiens* | Male |
| 9 | - | - | AUS 080 | Duckworth Laboratory, University of Cambridge | *Homo sapiens* | Male |
| 10 | - | - | AUS 081 | Duckworth Laboratory, University of Cambridge | *Homo sapiens* | Male |
| 11 | - | - | AUS 104 | Duckworth Laboratory, University of Cambridge | *Homo sapiens* | Male |
| 12 | - | - | AUS 106 | Duckworth Laboratory, University of Cambridge | *Homo sapiens* | Male |
| 13 | - | - | AUS 108 | Duckworth Laboratory, University of Cambridge | *Homo sapiens* | Male |
| 14 | - | - | AUS 122 | Duckworth Laboratory, University of Cambridge | *Homo sapiens* | Male |
| 15 | - | - | AUS 124 | Duckworth Laboratory, University of Cambridge | *Homo sapiens* | Female |
| 16 | - | - | AUS 125 | Duckworth Laboratory, University of Cambridge | *Homo sapiens* | Female |
| 17 | - | - | AUS 126 | Duckworth Laboratory, University of Cambridge | *Homo sapiens* | Female |
| 18 | - | - | AUS 127 | Duckworth Laboratory, University of Cambridge | *Homo sapiens* | Male |
| 19 | - | - | AUS 128 | Duckworth Laboratory, University of Cambridge | *Homo sapiens* | Male |
| 20 | - | - | AUS 129 | Duckworth Laboratory, University of Cambridge | *Homo sapiens* | Male |
| 21 | - | - | AUS 131 | Duckworth Laboratory, University of Cambridge | *Homo sapiens* | Female |
| 22 | 000102415 | MorphoSource | usnm:mammals:220380 | Smithsonian Institution National Museum of Natural History Collection | *Gorilla gorilla* | Female |
| 23 | 000102517 | MorphoSource | usnm:mammals:252581 | Smithsonian Institution National Museum of Natural History Collection | *Gorilla gorilla* | Female |
| 24 | 000102539 | MorphoSource | usnm:mammals:174720 | Smithsonian Institution National Museum of Natural History Collection | *Gorilla gorilla* | Male |
| 25 | 000102549 | MorphoSource | usnm:mammals:176213 | Smithsonian Institution National Museum of Natural History Collection | *Gorilla gorilla* | Male |
| 26 | 000102566 | MorphoSource | usnm:mammals:176210 | Smithsonian Institution National Museum of Natural History Collection | *Gorilla gorilla* | Male |
| 27 | 000102647 | MorphoSource | usnm:mammals:220324 | Smithsonian Institution National Museum of Natural History Collection | *Gorilla gorilla* | Male |
| 28 | 000102650 | MorphoSource | usnm:mammals:252576 | Smithsonian Institution National Museum of Natural History Collection | *Gorilla gorilla* | Female |
| 29 | 000102683 | MorphoSource | usnm:mammals:599167 | Smithsonian Institution National Museum of Natural History Collection | *Gorilla gorilla* | Male |
| 30 | 000102847 | MorphoSource | usnm:mammals:176215 | Smithsonian Institution National Museum of Natural History Collection | *Gorilla gorilla* | Male |
| 31 | 000102848 | MorphoSource | usnm:mammals:174717 | Smithsonian Institution National Museum of Natural History Collection | *Gorilla gorilla* | Male |
| 32 | 000102916 | MorphoSource | usnm:mammals:252579 | Smithsonian Institution National Museum of Natural History Collection | *Gorilla gorilla* | Female |
| 33 | 000102919 | MorphoSource | usnm:mammals:176207 | Smithsonian Institution National Museum of Natural History Collection | *Gorilla gorilla* | Male |
| 34 | 000102952 | MorphoSource | usnm:mammals:220060 | Smithsonian Institution National Museum of Natural History Collection | *Gorilla gorilla* | Female |
| 35 | 000102973 | MorphoSource | usnm:mammals:174722 | Smithsonian Institution National Museum of Natural History Collection | *Gorilla gorilla* | Male |
| 36 | 000103016 | MorphoSource | usnm:mammals:174712 | Smithsonian Institution National Museum of Natural History Collection | *Gorilla gorilla* | Male |
| 37 | 000103026 | MorphoSource | usnm:mammals:574138 | Smithsonian Institution National Museum of Natural History Collection | *Gorilla gorilla* | Male |
| 38 | 000103103 | MorphoSource | usnm:mammals:174716 | Smithsonian Institution National Museum of Natural History Collection | *Gorilla gorilla* | Male |
| 39 | 000103167 | MorphoSource | usnm:mammals:252577 | Smithsonian Institution National Museum of Natural History Collection | *Gorilla gorilla* | Female |

| | | | | | | |
|---|---|---|---|---|---|---|
| 40 | 000103194 | MorphoSource | usnm:mammals:220325 | Smithsonian Institution National Museum of Natural History Collection | *Gorilla gorilla* | Male |
| 41 | 000103213 | MorphoSource | usnm:mammals:297857 | Smithsonian Institution National Museum of Natural History Collection | *Gorilla gorilla* | Male |
| 42 | 000103224 | MorphoSource | usnm:mammals:599165 | Smithsonian Institution National Museum of Natural History Collection | *Gorilla gorilla* | Male |
| 43 | 000103296 | MorphoSource | usnm:mammals:176216 | Smithsonian Institution National Museum of Natural History Collection | *Gorilla gorilla* | Male |
| 44 | 000103349 | MorphoSource | usnm:mammals:176211 | Smithsonian Institution National Museum of Natural History Collection | *Gorilla gorilla* | Male |
| 45 | 000103348 | MorphoSource | usnm:mammals:176205 | Smithsonian Institution National Museum of Natural History Collection | *Gorilla gorilla* | Male |
| 46 | 000103357 | MorphoSource | usnm:mammals:176209 | Smithsonian Institution National Museum of Natural History Collection | *Gorilla gorilla* | Male |
| 47 | 000103399 | MorphoSource | usnm:mammals:174715 | Smithsonian Institution National Museum of Natural History Collection | *Gorilla gorilla* | Male |
| 48 | 000103429 | MorphoSource | usnm:mammals:599166 | Smithsonian Institution National Museum of Natural History Collection | *Gorilla gorilla* | Male |
| 49 | 000103463 | MorphoSource | usnm:mammals:582726 | Smithsonian Institution National Museum of Natural History Collection | *Gorilla gorilla* | Female |
| 50 | 000103487 | MorphoSource | usnm:mammals:197037 | Smithsonian Institution National Museum of Natural History Collection | *Gorilla gorilla* | Male |
| 51 | 000103499 | MorphoSource | usnm:mammals:174713 | Smithsonian Institution National Museum of Natural History Collection | *Gorilla gorilla* | Male |
| 52 | 000103562 | MorphoSource | usnm:mammals:154554 | Smithsonian Institution National Museum of Natural History Collection | *Gorilla gorilla* | Male |
| 53 | 000103652 | MorphoSource | usnm:mammals:252575 | Smithsonian Institution National Museum of Natural History Collection | *Gorilla gorilla* | Female |
| 54 | 000102414 | MorphoSource | usnm:mammals:396934 | Smithsonian Institution National Museum of Natural History Collection | *Gorilla beringei* | Male |
| 55 | 000102621 | MorphoSource | usnm:mammals:545030 | Smithsonian Institution National Museum of Natural History Collection | *Gorilla beringei* | Female |
| 56 | 000102627 | MorphoSource | usnm:mammals:396935 | Smithsonian Institution National Museum of Natural History Collection | *Gorilla beringei* | Female |
| 57 | 000102649 | MorphoSource | usnm:mammals:545032 | Smithsonian Institution National Museum of Natural History Collection | *Gorilla beringei* | Male |
| 58 | 000102720 | MorphoSource | usnm:mammals:545031 | Smithsonian Institution National Museum of Natural History Collection | *Gorilla beringei* | Female |
| 59 | 000102751 | MorphoSource | usnm:mammals:545036 | Smithsonian Institution National Museum of Natural History Collection | *Gorilla beringei* | Male |
| 60 | 000102762 | MorphoSource | usnm:mammals:239883 | Smithsonian Institution National Museum of Natural History Collection | *Gorilla beringei* | Male |
| 61 | 000102983 | MorphoSource | usnm:mammals:396942 | Smithsonian Institution National Museum of Natural History Collection | *Gorilla beringei* | Male |
| 62 | 000103006 | MorphoSource | usnm:mammals:397356 | Smithsonian Institution National Museum of Natural History Collection | *Gorilla beringei* | Female |
| 63 | 000103015 | MorphoSource | usnm:mammals:545035 | Smithsonian Institution National Museum of Natural History Collection | *Gorilla beringei* | Male |
| 64 | 000103038 | MorphoSource | usnm:mammals:545034 | Smithsonian Institution National Museum of Natural History Collection | *Gorilla beringei* | Male |
| 65 | 000103614 | MorphoSource | usnm:mammals:395636 | Smithsonian Institution National Museum of Natural History Collection | *Gorilla beringei* | Male |
| 66 | 000102486 | MorphoSource | usnm:mammals:395820 | Smithsonian Institution National Museum of Natural History Collection | *Pan troglodytes* | Male |
| 67 | 000102895 | MorphoSource | usnm:mammals:282763 | Smithsonian Institution National Museum of Natural History Collection | *Pan troglodytes* | Female |
| 68 | 000103302 | MorphoSource | usnm:mammals:84655 | Smithsonian Institution National Museum of Natural History Collection | *Pan troglodytes* | Female |
| 69 | 000102136 | MorphoSource | usnm:mammals:176235 | Smithsonian Institution National Museum of Natural History Collection | *Pan troglodytes* | Male |
| 70 | 000102182 | MorphoSource | usnm:mammals:176226 | Smithsonian Institution National Museum of Natural History Collection | *Pan troglodytes* | Male |
| 71 | 000102425 | MorphoSource | usnm:mammals:174710 | Smithsonian Institution National Museum of Natural History Collection | *Pan troglodytes* | Female |
| 72 | 000102434 | MorphoSource | usnm:mammals:174704 | Smithsonian Institution National Museum of Natural History Collection | *Pan troglodytes* | Male |
| 73 | 000102542 | MorphoSource | usnm:mammals:599172 | Smithsonian Institution National Museum of Natural History Collection | *Pan troglodytes* | Male |
| 74 | 000102558 | MorphoSource | usnm:mammals:599173 | Smithsonian Institution National Museum of Natural History Collection | *Pan troglodytes* | Female |
| 75 | 000102812 | MorphoSource | usnm:mammals:220065 | Smithsonian Institution National Museum of Natural History Collection | *Pan troglodytes* | Male |
| 76 | 000102885 | MorphoSource | usnm:mammals:174700 | Smithsonian Institution National Museum of Natural History Collection | *Pan troglodytes* | Female |
| 77 | 000102982 | MorphoSource | usnm:mammals:176228 | Smithsonian Institution National Museum of Natural History Collection | *Pan troglodytes* | Male |
| 78 | 000103008 | MorphoSource | usnm:mammals:220062 | Smithsonian Institution National Museum of Natural History Collection | *Pan troglodytes* | Female |
| 79 | 000103180 | MorphoSource | usnm:mammals:220064 | Smithsonian Institution National Museum of Natural History Collection | *Pan troglodytes* | Female |
| 80 | 000103246 | MorphoSource | usnm:mammals:220327 | Smithsonian Institution National Museum of Natural History Collection | *Pan troglodytes* | Male |
| 81 | 000103449 | MorphoSource | usnm:mammals:174701 | Smithsonian Institution National Museum of Natural History Collection | *Pan troglodytes* | Female |
| 82 | 000103498 | MorphoSource | usnm:mammals:220063 | Smithsonian Institution National Museum of Natural History Collection | *Pan troglodytes* | Female |

| | | | | | | |
|---|---|---|---|---|---|---|
| 83 | 000102438 | MorphoSource | usnm:mammals:143587 | Smithsonian Institution National Museum of Natural History Collection | *Pongo abelii* | Male |
| 84 | 000102533 | MorphoSource | usnm:mammals:143601 | Smithsonian Institution National Museum of Natural History Collection | *Pongo abelii* | Female |
| 85 | 000102544 | MorphoSource | usnm:mammals:143598 | Smithsonian Institution National Museum of Natural History Collection | *Pongo abelii* | Female |
| 86 | 000102578 | MorphoSource | usnm:mammals:143596 | Smithsonian Institution National Museum of Natural History Collection | *Pongo abelii* | Female |
| 87 | 000102743 | MorphoSource | usnm:mammals:143593 | Smithsonian Institution National Museum of Natural History Collection | *Pongo abelii* | Male |
| 88 | 000102871 | MorphoSource | usnm:mammals:143594 | Smithsonian Institution National Museum of Natural History Collection | *Pongo abelii* | Male |
| 89 | 000102884 | MorphoSource | usnm:mammals:143602 | Smithsonian Institution National Museum of Natural History Collection | *Pongo abelii* | Female |
| 90 | 000102939 | MorphoSource | usnm:mammals:267325 | Smithsonian Institution National Museum of Natural History Collection | *Pongo abelii* | Male |
| 91 | 000102959 | MorphoSource | usnm:mammals:270807 | Smithsonian Institution National Museum of Natural History Collection | *Pongo abelii* | Female |
| 92 | 000103300 | MorphoSource | usnm:mammals:143588 | Smithsonian Institution National Museum of Natural History Collection | *Pongo abelii* | Male |
| 93 | 000103378 | MorphoSource | usnm:mammals:578647 | Smithsonian Institution National Museum of Natural History Collection | *Pongo abelii* | Male |
| 94 | 000103595 | MorphoSource | usnm:mammals:143597 | Smithsonian Institution National Museum of Natural History Collection | *Pongo abelii* | Female |
| 95 | 000102426 | MorphoSource | usnm:mammals:153819 | Smithsonian Institution National Museum of Natural History Collection | *Pongo pygmaeus* | Female |
| 96 | 000102480 | MorphoSource | usnm:mammals:153828 | Smithsonian Institution National Museum of Natural History Collection | *Pongo pygmaeus* | Female |
| 97 | 000102604 | MorphoSource | usnm:mammals:142191 | Smithsonian Institution National Museum of Natural History Collection | *Pongo pygmaeus* | Female |
| 98 | 000102659 | MorphoSource | usnm:mammals:145309 | Smithsonian Institution National Museum of Natural History Collection | *Pongo pygmaeus* | Female |
| 99 | 000102727 | MorphoSource | usnm:mammals:145304 | Smithsonian Institution National Museum of Natural History Collection | *Pongo pygmaeus* | Male |
| 100 | 000102769 | MorphoSource | usnm:mammals:142196 | Smithsonian Institution National Museum of Natural History Collection | *Pongo pygmaeus* | Male |
| 101 | 000102799 | MorphoSource | usnm:mammals:153834 | Smithsonian Institution National Museum of Natural History Collection | *Pongo pygmaeus* | Male |
| 102 | 000102829 | MorphoSource | usnm:mammals:145321 | Smithsonian Institution National Museum of Natural History Collection | *Pongo pygmaeus* | Female |
| 103 | 000102869 | MorphoSource | usnm:mammals:153822 | Smithsonian Institution National Museum of Natural History Collection | *Pongo pygmaeus* | Female |
| 104 | 000103055 | MorphoSource | usnm:mammals:153805 | Smithsonian Institution National Museum of Natural History Collection | *Pongo pygmaeus* | Female |

### SOM Table S2

Landmark definitions

| No. | Name | Definition |
|---|---|---|
| 1 | M3 distal | Most distal point of the M3 socket |
| 2 | M2 buccal | Distobuccal corner of the M2 socket |
| 3 | M1 buccal | Distobuccal corner of the M1 socket |
| 4 | P4 buccal | Distobuccal corner of the P4 socket |
| 5 | P3 buccal | Distobuccal corner of the P3 socket |
| 6 | C buccal | Distobuccal corner of the C socket |
| 7 | I2 buccal | Distobuccal corner of the I2 socket |
| 8 | I2 lingual | Distolingual corner of the I2 socket |
| 9 | C lingual | Distolingual corner of the C socket |
| 10 | P3 lingual | Distolingual corner of the P3 socket |
| 11 | P4 lingual | Distolingual corner of the P4 socket |
| 12 | M1 lingual | Distolingual corner of the M1 socket |
| 13 | M2 lingual | Distolingual corner of the M2 socket |
| 14 | Infradentale | Midpoint between both central incisors at the lingual side |
| 15 | Linguale | Midpoint between both central incisors at the buccal side |
| 16 | Gnathion | Most inferior point at the symphysis |
| 17 | Condylar posterior | Most posterior point of the condylar articular surface |
| 18 | Condylar anterior | Most anterior point of the condylar articular surface |
| 19 | Coronion | Most superior point on the coronoid process |
| 20 | Extramolar sulcus | Most lateral point on the oblique line at the level of landmark 2 |
| 21 | Extramolar sulcus - Coronion (n= 20) | A curve along the oblique line and the anterior margin of the ramus (from landmark 19 to 20) |
| 22 | Mandibular notch (n= 10) | A curve along the mandibular notch (from landmark 18 to 19) |
| 23 | Gnathion - Condyle (n= 20) | A curve along the base of the mandible and the posterior margin of the ramus (from landmark 16 to 17) |

**SOM Table S3**

Definition of linear measurements extracted from landmarks

| # | Measurement | Landmark names | Landmark numbers |
|---|---|---|---|
| I | Symphysis height | Infradentale - Gnathion | 14 - 16 |
| II | Corpus width (from alveolar $M_2$-$M_3$) | M2 lingual - Extramolar sulcus | 13 - 20 |
| III | Coronoid projection | Gnathion - Coronion | 16 - 19 |
| IV | Condyle projection | Gnathion - Condylar posterior | 16 -17 |
| V | Mandibular notch | Coronion - Condylar posterior | 19 -17 |

### SOM Table S4A

Pairwise absolute differences in shape dimorphism

| | d | UCL (95%) | Z | *p-value* |
|---|---|---|---|---|
| *G. beringei : G. gorilla* | 5.00E-05 | 0.01410234 | -2.3132001 | 0.995 |
| *G. beringei : H. sapiens* | 1.59E-03 | 0.01440937 | -1.0248999 | 0.836 |
| *G. beringei : P. abelii* | 5.34E-03 | 0.01412082 | 0.171267 | 0.448 |
| *G. beringei : P. pygmaeus* | 1.55E-02 | 0.01592127 | 1.5738212 | 0.056 |
| *G. beringei : P. troglodytes* | 8.48E-04 | 0.01409871 | -1.3752822 | 0.897 |
| *G. gorilla : H. sapiens* | 1.64E-03 | 0.01058518 | -0.7119081 | 0.754 |
| *G. gorilla : P. abelii* | 5.29E-03 | 0.01391977 | 0.2085171 | 0.441 |
| *G. gorilla : P. pygmaeus* | 1.55E-02 | 0.01713369 | 1.3886913 | 0.087 |
| *G. gorilla : P. troglodytes* | 8.98E-04 | 0.01108934 | -1.2131754 | 0.872 |
| *H. sapiens : P. abelii* | 6.92E-03 | 0.01352081 | 0.5661286 | 0.285 |
| ***H. sapiens : P. pygmaeus*** | **1.71E-02** | **0.01648082** | **1.6654252** | **0.044** |
| *H. sapiens : P. troglodytes* | 7.41E-04 | 0.01139041 | -1.3340698 | 0.899 |
| *P. abelii : P. pygmaeus* | 1.02E-02 | 0.01560466 | 0.9222453 | 0.204 |
| *Pabelii : Ptroglodytes* | 6.18E-03 | 0.01348731 | 0.3809354 | 0.37 |
| *P. pygmaeus : P. troglodytes* | 1.64E-02 | 0.01670875 | 1.5891162 | 0.054 |

*Associated *p-values* <0.05 marked in bold

### SOM Table S4B

Pairwise absolute differences in size dimorphism

| | d | UCL (95%) | Z | *p-value* |
|---|---|---|---|---|
| *G. beringei : G. gorilla* | 0.046884658 | 0.2360747 | -0.92461569 | 0.7935 |
| *G. beringei : H. sapiens* | 0.172380699 | 0.2360747 | 0.81667572 | 0.2515 |
| *G. beringei : P. abelii* | 0.023589499 | 0.2360747 | -1.35420407 | 0.8055 |
| *G. beringei : P. pygmaeus* | 0.057029708 | 0.2360747 | -0.68027652 | 0.716 |
| *G. beringei : P. troglodytes* | 0.179045033 | 0.2360747 | 0.88589014 | 0.177 |
| *G. gorilla : H. sapiens* | 0.125496042 | 0.2360747 | 0.22613669 | 0.4945 |
| *G. gorilla : P. abelii* | 0.023295159 | 0.2360747 | -1.32676837 | 0.9135 |
| *G. gorilla : P. pygmaeus* | 0.103914366 | 0.2360747 | -0.04752352 | 0.5845 |
| *G. gorilla : P. troglodytes* | 0.132160375 | 0.2360747 | 0.37669344 | 0.426 |
| *H. sapiens : P. abelii* | 0.148791201 | 0.2360747 | 0.61304774 | 0.3525 |
| *H. sapiens : P. pygmaeus* | 0.229410408 | 0.2360747 | 1.40364907 | 0.1175 |
| *H. sapiens : P. troglodytes* | 0.006664333 | 0.2360747 | -1.86414805 | 0.992 |
| *P. abelii : P. pygmaeus* | 0.080619207 | 0.2360747 | -0.33756797 | 0.623 |
| *Pabelii : Ptroglodytes* | 0.155455534 | 0.2360747 | 0.63870691 | 0.321 |
| ***P. pygmaeus : P. troglodytes*** | **0.236074741** | **0.2360747** | **1.50823602** | **0.047** |

*Associated *p-values* <0.05 marked in bold

### SOM Table S5

Procrustes ANOVA and pairwise results for shape differences between species

| | Df | SS | F | *p-value* |
|---|---|---|---|---|
| **Inner corpus** | **5** | **0.17023** | **18.352** | **0.001** |
| **Outer corpus** | **5** | **0.44466** | **39.759** | **0.001** |
| **Inner ramus** | **5** | **0.15808** | **13.812** | **0.001** |
| **Outer ramus** | **5** | **0.081721** | **12.878** | **0.001** |

| Inner corpus | d | UCL (95%) | Z | *p-value* |
|---|---|---|---|---|
| ***G. beringei : G. gorilla*** | **4.91E-04** | **2.30E-04** | **3.1149262** | **0.001** |
| ***G. beringei : H. sapiens*** | **7.56E-04** | **2.03E-04** | **3.9406625** | **0.001** |
| ***G. beringei : P. abelii*** | **6.04E-04** | **2.14E-04** | **3.4028069** | **0.001** |
| ***G. beringei : P. pygmaeus*** | **3.77E-04** | **2.17E-04** | **2.4531849** | **0.005** |
| ***G. beringei : P. troglodytes*** | **5.62E-04** | **1.98E-04** | **3.3386281** | **0.001** |
| ***G. gorilla : H. sapiens*** | **1.25E-03** | **9.52E-05** | **7.1801894** | **0.001** |
| *G. gorilla : P. abelii* | 1.12E-04 | 2.31E-04 | 0.5449132 | 0.285 |
| ***G. gorilla : P. pygmaeus*** | **8.68E-04** | **2.67E-04** | **4.364172** | **0.001** |
| *G. gorilla : P. troglodytes* | 7.04E-05 | 1.31E-04 | 0.6853099 | 0.239 |
| ***H. sapiens : P. abelii*** | **1.36E-03** | **2.14E-04** | **5.1004417** | **0.001** |
| ***H. sapiens : P. pygmaeus*** | **3.80E-04** | **2.41E-04** | **2.3751875** | **0.012** |
| ***H. sapiens : P. troglodytes*** | **1.32E-03** | **1.23E-04** | **5.8155442** | **0.001** |
| ***P. abelii : P. pygmaeus*** | **9.80E-04** | **2.33E-04** | **4.2087713** | **0.001** |
| *Pabelii : Ptroglodytes* | 4.21E-05 | 2.05E-04 | -0.1484959 | 0.553 |
| ***P. pygmaeus : P. troglodytes*** | **9.38E-04** | **2.24E-04** | **3.9285299** | **0.001** |

| Outer corpus | d | UCL (95%) | Z | *p-value* |
|---|---|---|---|---|
| ***G. beringei : G. gorilla*** | **0.000542196** | **0.000557696** | **1.64166622** | **0.054** |
| ***G. beringei : H. sapiens*** | **0.005818066** | **0.000534365** | **4.97607317** | **0.001** |
| ***G. beringei : P. abelii*** | **0.001017734** | **0.000658891** | **2.18094917** | **0.008** |
| ***G. beringei : P. pygmaeus*** | **0.000868331** | **0.000686145** | **1.89800936** | **0.028** |
| ***G. beringei : P. troglodytes*** | **0.000736303** | **0.000515198** | **2.02052322** | **0.021** |
| ***G. gorilla : H. sapiens*** | **0.006360262** | **0.000285374** | **6.60503** | **0.001** |
| *G. gorilla : P. abelii* | 0.000475538 | 0.000661899 | 1.36225686 | 0.11 |
| *G. gorilla : P. pygmaeus* | 0.000326135 | 0.000633603 | 0.83155028 | 0.207 |
| *G. gorilla : P. troglodytes* | 0.000194106 | 0.00036661 | 0.94134161 | 0.161 |
| ***H. sapiens : P. abelii*** | **0.006835801** | **0.00063951** | **5.04195712** | **0.001** |
| ***H. sapiens : P. pygmaeus*** | **0.006686397** | **0.000592378** | **4.99874722** | **0.001** |
| ***H. sapiens : P. troglodytes*** | **0.006554369** | **0.00035587** | **6.04218727** | **0.001** |
| *P. abelii : P. pygmaeus* | 0.000149403 | 0.000737614 | 0.05057193 | 0.474 |

| | | | | |
|---|---|---|---|---|
| *Pabelii : Ptroglodytes* | 0.000281432 | 0.000653965 | 0.84184841 | 0.207 |
| *P. pygmaeus : P. troglodytes* | 0.000132029 | 0.000605949 | 0.08643108 | 0.468 |

| **Inner ramus** | **d** | **UCL (95%)** | **Z** | ***p-value*** |
|---|---|---|---|---|
| ***G. beringei : G. gorilla*** | **7.11E-04** | **0.000231592** | **4.1414749** | **0.001** |
| ***G. beringei : H. sapiens*** | **1.03E-03** | **0.000209843** | **4.5571628** | **0.001** |
| *G. beringei : P. abelii* | 1.22E-04 | 0.000213845 | 0.9169376 | 0.182 |
| *G. beringei : P. pygmaeus* | 1.20E-04 | 0.00022464 | 0.7917029 | 0.224 |
| ***G. beringei : P. troglodytes*** | **3.11E-04** | **0.000188318** | **2.3272263** | **0.011** |
| ***G. gorilla : H. sapiens*** | **1.74E-03** | **0.000102542** | **8.151448** | **0.001** |
| ***G. gorilla : P. abelii*** | **5.89E-04** | **0.000245673** | **3.6952321** | **0.001** |
| ***G. gorilla : P. pygmaeus*** | **5.91E-04** | **0.000270529** | **3.6812875** | **0.002** |
| ***G. gorilla : P. troglodytes*** | **4.00E-04** | **0.000142759** | **3.5487365** | **0.002** |
| ***H. sapiens : P. abelii*** | **1.15E-03** | **0.000209413** | **5.1330495** | **0.001** |
| ***H. sapiens : P. pygmaeus*** | **1.15E-03** | **0.000256699** | **5.2540581** | **0.001** |
| ***H. sapiens : P. troglodytes*** | **1.34E-03** | **0.00013586** | **5.8249218** | **0.001** |
| *P. abelii : P. pygmaeus* | 2.49E-06 | 0.000224325 | -2.1051767 | 0.98 |
| *Pabelii : Ptroglodytes* | 1.89E-04 | 0.000213791 | 1.5455994 | 0.066 |
| *P. pygmaeus : P. troglodytes* | 1.91E-04 | 0.000220543 | 1.3167469 | 0.082 |

| **Outer ramus** | **d** | **UCL (95%)** | **Z** | ***p-value*** |
|---|---|---|---|---|
| ***G. beringei : G. gorilla*** | **3.10E-04** | **1.27E-04** | **3.6425281** | **0.001** |
| ***G. beringei : H. sapiens*** | **3.09E-04** | **1.10E-04** | **3.3677203** | **0.001** |
| ***G. beringei : P. abelii*** | **2.83E-04** | **1.07E-04** | **3.2504051** | **0.001** |
| *G. beringei : P. pygmaeus* | 2.78E-05 | 1.32E-04 | -0.1197258 | 0.55 |
| ***G. beringei : P. troglodytes*** | **1.68E-04** | **1.09E-04** | **2.3456493** | **0.011** |
| ***G. gorilla : H. sapiens*** | **6.19E-04** | **5.47E-05** | **7.1828836** | **0.001** |
| ***G. gorilla : P. abelii*** | **5.93E-04** | **1.20E-04** | **5.9215788** | **0.001** |
| ***G. gorilla : P. pygmaeus*** | **3.38E-04** | **1.63E-04** | **3.4351913** | **0.001** |
| ***G. gorilla : P. troglodytes*** | **1.42E-04** | **7.82E-05** | **2.7492625** | **0.005** |
| *H. sapiens : P. abelii* | 2.58E-05 | 1.06E-04 | -0.1731852 | 0.562 |
| ***H. sapiens : P. pygmaeus*** | **2.81E-04** | **1.43E-04** | **2.9266129** | **0.003** |
| ***H. sapiens : P. troglodytes*** | **4.77E-04** | **6.73E-05** | **5.4150876** | **0.001** |
| ***P. abelii : P. pygmaeus*** | **2.55E-04** | **1.27E-04** | **2.8940551** | **0.002** |
| ***Pabelii : Ptroglodytes*** | **4.51E-04** | **1.02E-04** | **4.3914912** | **0.001** |
| ***P. pygmaeus : P. troglodytes*** | **1.96E-04** | **1.37E-04** | **2.2420216** | **0.012** |

*Associated *p-values* <0.05 marked in bold

### SOM Table S6

Procrustes variances and corresponding post-hoc pairwise differences between Procrustes variances

| Species | Inner corpus | Outer corpus | Inner ramus | Outer ramus |
|---|---|---|---|---|
| *G. beringei* | 0.001254239 | 0.001281835 | 0.001386702 | 0.000727669 |
| *G. gorilla* | 0.001968352 | 0.001692238 | 0.001697622 | 0.00091153 |
| *P. pygmaeus* | 0.001172679 | 0.001315654 | 0.00168529 | 0.000924963 |
| *P. abelii* | 0.002067617 | 0.00207681 | 0.002335662 | 0.00136833 |
| *P. troglodytes* | 0.001158627 | 0.001433538 | 0.002380246 | 0.001367977 |
| *H. sapiens* | 0.002263296 | 0.004153465 | 0.003239291 | 0.001788127 |

| **Inner corpus** | Pairwise differences | *p-value* |
|---|---|---|
| *G. beringei : G. gorilla* | $7.14^{-04}$ | 0.088 |
| ***G. beringei : H. sapiens*** | **$1.01^{-03}$** | **0.032** |
| *G. beringei : P. abelii* | $8.13^{-04}$ | 0.124 |
| *G. beringei : P. pygmaeus* | $8.16^{-05}$ | 0.869 |
| *G. beringei : P. troglodytes* | $9.56^{-05}$ | 0.834 |
| *G. gorilla : H. sapiens* | $2.95^{-04}$ | 0.444 |
| *G. gorilla : P. abelii* | $9.93^{-05}$ | 0.815 |
| *G. gorilla : P. pygmaeus* | $7.96^{-04}$ | 0.095 |
| *G. gorilla : P. troglodytes* | $8.10^{-04}$ | **0.032** |
| *H. sapiens : P. abelii* | 0.000195679 | 0.67 |
| ***H. sapiens : P. pygmaeus*** | **0.001090618** | **0.025** |
| ***H. sapiens : P. troglodytes*** | **0.001104669** | **0.008** |
| *P. abelii : P. pygmaeus* | $8.95^{-04}$ | 0.109 |
| *Pabelii : Ptroglodytes* | $9.09^{-04}$ | 0.055 |
| *P. pygmaeus : P. troglodytes* | $1.41^{-05}$ | 0.973 |

| **Outer corpus** | Pairwise differences | *p-value* |
|---|---|---|
| *G. beringei : G. gorilla* | $4.10^{-04}$ | 0.489 |
| ***G. beringei : H. sapiens*** | **$2.87^{-03}$** | **0.001** |
| *G. beringei : P. abelii* | $7.95^{-04}$ | 0.279 |
| *G. beringei : P. pygmaeus* | $3.38^{-05}$ | 0.966 |
| *G. beringei : P. troglodytes* | $1.52^{-04}$ | 0.835 |
| ***G. gorilla : H. sapiens*** | **0.002461226** | **0.001** |
| *G. gorilla : P. abelii* | 0.000384572 | 0.482 |
| *G. gorilla : P. pygmaeus* | 0.000376584 | 0.581 |
| *G. gorilla : P. troglodytes* | 0.0002587 | 0.638 |
| ***H. sapiens : P. abelii*** | **0.002076655** | **0.004** |
| ***H. sapiens : P. pygmaeus*** | **0.002837811** | **0.001** |
| ***H. sapiens : P. troglodytes*** | **0.002719927** | **0.001** |
| *P. abelii : P. pygmaeus* | 0.000761156 | 0.302 |
| *Pabelii : Ptroglodytes* | 0.000643272 | 0.314 |
| *P. pygmaeus : P. troglodytes* | $1.18^{-04}$ | 0.854 |

| **Inner ramus** | Pairwise differences | *p-value* |
|---|---|---|
| *G. beringei : G. gorilla* | 0.00031092 | 0.404 |
| ***G. beringei : H. sapiens*** | **0.001852589** | **0.001** |
| ***G. beringei : P. abelii*** | **0.00094896** | **0.035** |
| *G. beringei : P. pygmaeus* | 0.000298588 | 0.547 |
| ***G. beringei : P. troglodytes*** | **0.000993544** | **0.014** |
| ***G. gorilla : H. sapiens*** | **$1.54^{-03}$** | **0.001** |
| *G. gorilla : P. abelii* | $6.38^{-04}$ | 0.101 |
| *G. gorilla : P. pygmaeus* | $1.23^{-05}$ | 0.973 |
| ***G. gorilla : P. troglodytes*** | **$6.83^{-04}$** | **0.042** |
| ***H. sapiens : P. abelii*** | **0.000903628** | **0.033** |
| ***H. sapiens : P. pygmaeus*** | **0.001554001** | **0.001** |
| ***H. sapiens : P. troglodytes*** | **0.000859045** | **0.022** |
| *P. abelii : P. pygmaeus* | $6.50^{-04}$ | 0.17 |
| *Pabelii : Ptroglodytes* | $4.46^{-05}$ | 0.916 |
| *P. pygmaeus : P. troglodytes* | $6.95^{-04}$ | 0.13 |

| **Outer ramus** | Pairwise differences | *p-value* |
|---|---|---|
| *G. beringei : G. gorilla* | 0.000183861 | 0.504 |
| ***G. beringei : H. sapiens*** | **0.001060458** | **0.001** |
| ***G. beringei : P. abelii*** | **0.00064066** | **0.044** |
| *G. beringei : P. pygmaeus* | 0.000197294 | 0.571 |
| ***G. beringei : P. troglodytes*** | **0.000640307** | **0.026** |
| ***G. gorilla : H. sapiens*** | **$8.77^{-04}$** | **0.001** |
| *G. gorilla : P. abelii* | $4.57^{-04}$ | 0.099 |
| *G. gorilla : P. pygmaeus* | $1.34^{-05}$ | 0.966 |
| *G. gorilla : P. troglodytes* | $4.56^{-04}$ | 0.062 |
| *H. sapiens : P. abelii* | 0.000419798 | 0.168 |
| ***H. sapiens : P. pygmaeus*** | **0.000863164** | **0.008** |
| *H. sapiens : P. troglodytes* | 0.00042015 | 0.105 |
| *P. abelii : P. pygmaeus* | $4.43^{-04}$ | 0.215 |
| *Pabelii : Ptroglodytes* | $3.53^{-07}$ | 1 |
| *P. pygmaeus : P. troglodytes* | $4.43^{-04}$ | 0.173 |

*Associated *p-values* <0.05 marked in bol

### SOM Table S7

Pairwise absolute differences in path distances of trajectory analyses

| *G. beringei : G. gorilla* | d | UCL (95%) | Z | *p-value* |
|---|---|---|---|---|
| Inner corpus | 0.012796655 | 0.01940667 | 0.91144469 | 0.191 |
| Outer corpus | 0.004479335 | 0.02003625 | -0.3520842 | 0.643 |
| Inner ramus | 0.00188341 | 0.01715576 | -1.0326673 | 0.824 |
| Outer ramus | 0.000571708 | 0.01557222 | -1.6406604 | 0.934 |
| *Gberingei:Hsapiens* | d | UCL (95%) | Z | *p-value* |
| Inner corpus | 0.017982176 | 0.01910759 | 1.52777426 | 0.061 |
| Outer corpus | 0.0030934 | 0.01856049 | -0.6567869 | 0.73 |
| Inner ramus | 0.002914729 | 0.01695915 | -0.6083943 | 0.713 |
| Outer ramus | 0.012458624 | 0.01496848 | 1.3158001 | 0.098 |
| *Gberingei:Pabelii* | d | UCL (95%) | Z | *p-value* |
| Inner corpus | 0.013548032 | 0.019205 | 1.09456857 | 0.148 |
| Outer corpus | 0.010645513 | 0.01758398 | 0.7330956 | 0.241 |
| Inner ramus | 0.005127587 | 0.01554189 | 0.0275376 | 0.509 |
| Outer ramus | 0.010155225 | 0.01405418 | 1.0568304 | 0.15 |
| *Gberingei:Ppygmaeus* | d | UCL (95%) | Z | *p-value* |
| Inner corpus | 0.008510612 | 0.02011028 | 0.39222301 | 0.364 |
| Outer corpus | 0.000373162 | 0.01958267 | -1.8820221 | 0.967 |
| **Inner ramus** | **0.020729613** | **0.01746836** | **1.902292** | **0.021** |
| Outer ramus | 0.00851372 | 0.01677089 | 0.6550107 | 0.27 |
| *Gberingei:Ptroglodytes* | d | UCL (95%) | Z | *p-value* |
| Inner corpus | 0.015933122 | 0.01947015 | 1.31405484 | 0.101 |
| Outer corpus | 0.005341848 | 0.018412 | -0.1096223 | 0.545 |
| Inner ramus | 0.003459729 | 0.01600098 | -0.3869862 | 0.644 |
| Outer ramus | 0.00042577 | 0.01530653 | -1.6443717 | 0.942 |
| *Ggorilla:Hsapiens* | d | UCL (95%) | Z | *p-value* |
| Inner corpus | 0.005185521 | 0.01525136 | 0.13993674 | 0.442 |
| Outer corpus | 0.001385935 | 0.01361465 | -1.0125905 | 0.841 |
| Inner ramus | 0.004798138 | 0.01179434 | 0.2435381 | 0.42 |
| **Outer ramus** | **0.011886916** | **0.01009286** | **1.8607483** | **0.023** |
| *Ggorilla:Pabelii* | d | UCL (95%) | Z | *p-value* |
| Inner corpus | 0.000751377 | 0.01815537 | -1.42098143 | 0.906 |
| Outer corpus | 0.015124848 | 0.01735578 | 1.3497399 | 0.089 |
| Inner ramus | 0.007010996 | 0.01621199 | 0.3200725 | 0.388 |
| Outer ramus | 0.009583517 | 0.01415274 | 0.9766373 | 0.178 |
| *Ggorilla:Ppygmaeus* | d | UCL (95%) | Z | *p-value* |
| Inner corpus | 0.004286044 | 0.02282794 | -0.56318037 | 0.711 |
| Outer corpus | 0.004852497 | 0.02191026 | -0.5366405 | 0.705 |
| **Inner ramus** | **0.022613022** | **0.02099308** | **1.7240372** | **0.031** |
| Outer ramus | 0.007942012 | 0.01859074 | 0.3029504 | 0.4 |

| *Ggorilla:Ptroglodytes* | d | UCL (95%) | Z | *p-value* |
|---|---|---|---|---|
| Inner corpus | 0.003136467 | 0.01576679 | -0.45955734 | 0.675 |
| Outer corpus | 0.000862513 | 0.01479264 | -1.4274047 | 0.914 |
| Inner ramus | 0.00157632 | 0.01210403 | -0.9227968 | 0.805 |
| Outer ramus | 0.000145938 | 0.01139835 | -2.0231835 | 0.976 |
| *Hsapiens:Pabelii* | d | UCL (95%) | Z | *p-value* |
| Inner corpus | 0.004434144 | 0.01818678 | -0.21032115 | 0.589 |
| Outer corpus | 0.013738912 | 0.01681148 | 1.2524813 | 0.1 |
| Inner ramus | 0.002212858 | 0.01582629 | -0.7755452 | 0.767 |
| Outer ramus | 0.002303399 | 0.01372161 | -0.5837967 | 0.722 |
| *Hsapiens:Ppygmaeus* | d | UCL (95%) | Z | *p-value* |
| Inner corpus | 0.009471564 | 0.02251013 | 0.36126801 | 0.362 |
| Outer corpus | 0.003466561 | 0.02105254 | -0.7581274 | 0.755 |
| Inner ramus | 0.017814884 | 0.0200769 | 1.3726173 | 0.085 |
| Outer ramus | 0.003944904 | 0.01803936 | -0.314264 | 0.609 |
| *Hsapiens:Ptroglodytes* | d | UCL (95%) | Z | *p-value* |
| Inner corpus | 0.002049054 | 0.01578338 | -0.81600978 | 0.783 |
| Outer corpus | 0.002248449 | 0.01428232 | -0.713172 | 0.752 |
| Inner ramus | 0.006374458 | 0.01178506 | 0.5577478 | 0.323 |
| **Outer ramus** | **0.012032854** | **0.01072638** | **1.7993652** | **0.028** |
| *Pabelii:Ppygmaeus* | d | UCL (95%) | Z | *p-value* |
| Inner corpus | 0.00503742 | 0.02047383 | -0.26405484 | 0.614 |
| Outer corpus | 0.010272351 | 0.0191466 | 0.6100053 | 0.289 |
| Inner ramus | 0.015602026 | 0.01713642 | 1.4170697 | 0.078 |
| Outer ramus | 0.001641505 | 0.01607052 | -0.9822552 | 0.823 |
| *Pabelii:Ptroglodytes* | d | UCL (95%) | Z | *p-value* |
| Inner corpus | 0.00238509 | 0.01849585 | -0.75996597 | 0.78 |
| Outer corpus | 0.015987361 | 0.01615255 | 1.5818569 | 0.052 |
| Inner ramus | 0.008587316 | 0.01516306 | 0.7271372 | 0.247 |
| Outer ramus | 0.009729455 | 0.01314315 | 1.0878582 | 0.14 |
| *Ppygmaeus:Ptroglodytes* | d | UCL (95%) | Z | *p-value* |
| Inner corpus | 0.00742251 | 0.02162225 | 0.08954791 | 0.475 |
| Outer corpus | 0.00571501 | 0.02090119 | -0.1912235 | 0.592 |
| **Inner ramus** | **0.024189342** | **0.01881799** | **2.032358** | **0.009** |
| Outer ramus | 0.00808795 | 0.01721141 | 0.4774653 | 0.323 |

*Associated *p-values* <0.05 marked in bold

### SOM Table S8

Pairwise comparisons between trajectory directions

| *G. beringei : G. gorilla* | r | angle | UCL (95%) | Z | *p-value* |
|---|---|---|---|---|---|
| Inner corpus | 0.55315229 | 0.9846529 | 1.678047 | -0.49204124 | 0.68 |
| Outer corpus | 0.49819005 | 1.0492862 | 1.406612 | 0.34322984 | 0.37 |
| Inner ramus | 0.25895823 | 1.3088528 | 1.616996 | 0.59805784 | 0.275 |
| Outer ramus | 0.33645594 | 1.227645 | 1.698649 | 0.21448443 | 0.432 |
| *Gberingei:Hsapiens* | r | angle | UCL (95%) | Z | *p-value* |
| **Inner corpus** | **-0.29460819** | **1.8698418** | **1.74651** | **1.92543116** | **0.023** |
| **Outer corpus** | **0.07578** | **1.4949436** | **1.45715** | **1.75971389** | **0.037** |
| Inner ramus | 0.02635228 | 1.544441 | 1.638045 | 1.33573786 | 0.094 |
| **Outer ramus** | **-0.32285428** | **1.89954** | **1.745193** | **2.08081876** | **0.016** |
| *Gberingei:Pabelii* | r | angle | UCL (95%) | Z | *p-value* |
| Inner corpus | -0.1814181 | 1.7532246 | 1.780454 | 1.55947255 | 0.062 |
| Outer corpus | 0.05887878 | 1.5118835 | 1.531945 | 1.62918799 | 0.055 |
| Inner ramus | 0.38631139 | 1.1741672 | 1.69644 | -0.217877 | 0.579 |
| Outer ramus | 0.21691106 | 1.352147 | 1.760773 | 0.39401533 | 0.355 |
| *Gberingei:Ppygmaeus* | r | angle | UCL (95%) | Z | *p-value* |
| Inner corpus | 0.32787125 | 1.2367469 | 1.823196 | -0.08468553 | 0.526 |
| Outer corpus | 0.37174775 | 1.1899053 | 1.568495 | 0.34630174 | 0.373 |
| Inner ramus | 0.30104589 | 1.2650071 | 1.734507 | 0.03159144 | 0.502 |
| Outer ramus | 0.07914917 | 1.491564 | 1.842224 | 0.64978268 | 0.267 |
| *Gberingei:Ptroglodytes* | r | angle | UCL (95%) | Z | *p-value* |
| Inner corpus | 0.56143759 | 0.9746743 | 1.744705 | -0.64951564 | 0.712 |
| Outer corpus | 0.6053907 | 0.9205397 | 1.443114 | -0.41665914 | 0.657 |
| Inner ramus | 0.24969723 | 1.3184288 | 1.636431 | 0.52834375 | 0.297 |
| Outer ramus | 0.37111166 | 1.19059 | 1.731855 | -0.01742551 | 0.508 |
| *Ggorilla:Hsapiens* | r | angle | UCL (95%) | Z | *p-value* |
| **Inner corpus** | **-0.21786067** | **1.7904183** | **1.584105** | **2.20606979** | **0.011** |
| **Outer corpus** | **0.19467324** | **1.374872** | **1.241396** | **2.01247061** | **0.02** |
| Inner ramus | 0.19307957 | 1.3764965 | 1.476189 | 1.33044523 | 0.105 |
| Outer ramus | 0.26029319 | 1.30747 | 1.544182 | 0.90815762 | 0.173 |
| *Ggorilla:Pabelii* | r | angle | UCL (95%) | Z | *p-value* |
| Inner corpus | 0.14711183 | 1.4231486 | 1.705251 | 0.9834328 | 0.164 |
| Outer corpus | 0.60401229 | 0.9222704 | 1.347688 | -0.0736633 | 0.528 |
| Inner ramus | 0.22218891 | 1.3467374 | 1.575333 | 0.86460344 | 0.197 |
| Outer ramus | 0.45117266 | 1.102717 | 1.668733 | -0.13224216 | 0.549 |
| *Ggorilla:Ppygmaeus* | r | angle | UCL (95%) | Z | *p-value* |
| Inner corpus | 0.19210668 | 1.377488 | 1.732791 | 0.67483957 | 0.253 |
| Outer corpus | 0.38006857 | 1.1809259 | 1.446126 | 0.74461109 | 0.245 |
| Inner ramus | 0.33087952 | 1.2335609 | 1.620394 | 0.27499971 | 0.393 |
| Outer ramus | 0.47295493 | 1.078155 | 1.717471 | -0.30775577 | 0.609 |
| *Ggorilla:Ptroglodytes* | r | angle | UCL (95%) | Z | *p-value* |

| | r | angle | UCL (95%) | Z | *p-value* |
|---|---|---|---|---|---|
| Inner corpus | 0.67808707 | 0.8256395 | 1.573073 | -0.84193152 | 0.789 |
| Outer corpus | 0.71188793 | 0.7786135 | 1.3026 | -0.51554843 | 0.702 |
| Inner ramus | 0.21653815 | 1.3525292 | 1.489891 | 1.11089487 | 0.14 |
| Outer ramus | 0.01492706 | 1.555869 | 1.585749 | 1.52058832 | 0.064 |
| *Hsapiens:Pabelii* | r | angle | UCL (95%) | Z | *p-value* |
| Inner corpus | 0.03940829 | 1.5313778 | 1.729932 | 1.12995857 | 0.141 |
| Outer corpus | 0.19326855 | 1.3763039 | 1.409647 | 1.56912935 | 0.063 |
| **Inner ramus** | **-0.26699282** | **1.8410675** | **1.62039** | **2.3668746** | **0.009** |
| **Outer ramus** | **-0.24469678** | **1.818003** | **1.698132** | **1.95800733** | **0.021** |
| *Hsapiens:Ppygmaeus* | r | angle | UCL (95%) | Z | *p-value* |
| Inner corpus | 0.23783796 | 1.330657 | 1.778811 | 0.43524817 | 0.349 |
| Outer corpus | 0.4402811 | 1.1148846 | 1.472079 | 0.36523339 | 0.372 |
| **Inner ramus** | **-0.15458713** | **1.7260059** | **1.658399** | **1.89135288** | **0.028** |
| Outer ramus | 0.18757111 | 1.382108 | 1.748083 | 0.62602218 | 0.273 |
| *Hsapiens:Ptroglodytes* | r | angle | UCL (95%) | Z | *p-value* |
| **Inner corpus** | **-0.30036546** | **1.8758721** | **1.636881** | **2.24420444** | **0.008** |
| **Outer corpus** | **0.02141746** | **1.5493772** | **1.335462** | **2.27795242** | **0.01** |
| **Inner ramus** | **-0.28476015** | **1.8595525** | **1.524922** | **2.6747068** | **0.003** |
| **Outer ramus** | **-0.73912227** | **2.402563** | **1.632173** | **3.33055396** | **0.001** |
| *Pabelii:Ppygmaeus* | r | angle | UCL (95%) | Z | *p-value* |
| Inner corpus | 0.19590504 | 1.373616 | 1.816697 | 0.39869531 | 0.356 |
| Outer corpus | 0.06390406 | 1.5068487 | 1.531723 | 1.54816982 | 0.061 |
| Inner ramus | 0.58234797 | 0.9491824 | 1.715395 | -1.23017806 | 0.884 |
| Outer ramus | 0.52303136 | 1.020393 | 1.821464 | -0.76578609 | 0.761 |
| *Pabelii:Ptroglodytes* | r | angle | UCL (95%) | Z | *p-value* |
| Inner corpus | 0.18862086 | 1.3810387 | 1.706574 | 0.75332413 | 0.24 |
| Outer corpus | 0.55121547 | 0.986976 | 1.383974 | 0.03760981 | 0.492 |
| Inner ramus | 0.46623802 | 1.0857628 | 1.565294 | -0.32253036 | 0.634 |
| Outer ramus | 0.34953895 | 1.213717 | 1.689146 | 0.24429199 | 0.411 |
| *Ppygmaeus:Ptroglodytes* | r | angle | UCL (95%) | Z | *p-value* |
| Inner corpus | 0.40123575 | 1.1579308 | 1.76361 | -0.08241095 | 0.536 |
| Outer corpus | 0.17332366 | 1.3965929 | 1.496349 | 1.34366044 | 0.104 |
| Inner ramus | 0.48225697 | 1.0675671 | 1.63315 | -0.53215091 | 0.699 |
| Outer ramus | 0.22622437 | 1.342597 | 1.732502 | 0.50621212 | 0.312 |

*Associated *p-values* <0.05 marked in bold
